# Multi-domain cognitive assessment of male mice reveals whole body exposure to space radiation is not detrimental to high-level cognition and actually improves pattern separation

**DOI:** 10.1101/796938

**Authors:** Cody W. Whoolery, Sanghee Yun, Ryan P. Reynolds, Melanie J. Lucero, Ivan Soler, Fionya H. Tran, Naoki Ito, Rachel L. Redfield, Devon R. Richardson, Hung-ying Shih, Phillip D. Rivera, Benjamin P. C. Chen, Shari G. Birnbaum, Ann M. Stowe, Amelia J. Eisch

## Abstract

Astronauts on interplanetary space missions - such as to Mars - will be exposed to space radiation, a spectrum of highly-charged, fast-moving particles that includes ^56^Fe and ^28^Si. Earth-based preclinical studies with mature, “astronaut-aged” rodents show space radiation decreases performance in low- and some high-level cognitive tasks. Given the prevalence of touchscreens in astronaut training and in-mission assessment, and the ability of rodent touchscreen tasks to assess the functional integrity of brain circuits and multiple cognitive domains in a non-aversive way, it is surprising the effect of space radiation on rodent touchscreen performance is unknown. To fill this knowledge gap, 6-month-old C57BL/6J male mice were exposed to whole-body space radiation and assessed on a touchscreen battery starting 1-month later. Relative to Sham, ^56^Fe irradiation did not overtly change performance on tasks of visual discrimination, reversal learning, rule-based, or object-spatial paired associates learning, suggesting preserved functional integrity of supporting brain circuits. Surprisingly, ^56^Fe irradiation led to better performance on a dentate gyrus-reliant task of pattern separation ability. Irradiated mice discriminated similar visual cues in ∼40% fewer days and ∼40% more accurately than control mice. Improved pattern separation was not touchscreen-, radiation-particle, or neurogenesis-dependent, as both ^56^Fe and ^28^Si irradiation led to faster context discrimination (e.g. Sham Block 5 vs. ^56^Fe Block 2) in a non-touchscreen task and ^56^Fe led to fewer new dentate gyrus neurons relative to Sham. These data urge revisitation of the broadly-held view that space radiation is detrimental to cognition.

**SIGNIFICANCE STATEMENT:** Astronauts on an interplanetary mission - such as to Mars - will be unavoidably exposed to galactic cosmic radiation, a spectrum of highly-charged, fast-moving particles. Rodent studies suggest space radiation is detrimental to cognition. However, here we show this is not universally true. Mature mice that received whole body exposure to Mars-relevant space radiation perform similarly to control mice on high-level cognitive tasks, reflecting the functional integrity of key neural circuits. Even more surprisingly, irradiated mice perform better than controls in both appetitive and aversive tests of pattern separation, a mission-critical task reliant on dentate gyrus integrity. Notably, improved pattern separation was not touchscreen-, radiation-particle-, or neurogenesis-dependent. Our work urges revisitation of the generally-accepted conclusion that space radiation is detrimental to cognition.

## Introduction

Interplanetary missions - such as to Mars - are a high priority for many national space agencies. The crew of future missions will face hazards to human health (1–6). Among these hazards is exposure to galactic cosmic radiation (7–12) which includes a spectrum of high-(H) atomic number (Z) and high-energy (E) particles such as ^56^Fe and ^28^Si. Fast-moving HZE particles cannot be effectively blocked by modern spacecraft shielding (13–20). Given their unavoidable nature, it is concerning that studies exposing laboratory animals to Earth-based space radiation generally conclude HZE particles are detrimental to brain and behavior (21–24). Such preclinical data suggest HZE particle exposure may be harmful to astronaut health and cognition and thus impede mission success.

However, there are several reasons to revisit the conclusion that HZE particle exposure is detrimental to cognitive function. First, age at the time of irradiation matters. Most preclinical data that led to the Probabilistic Risk Assessment of HZE particles being detrimental to cognition and related behavior were from tests performed on young adult rodents (∼2-3 months [mon] old at exposure)(23, 25); in many cases, age of the animals tested was not even reported (23). To more accurately reflect the average age of astronauts, NASA now requires ground-based space studies to be performed in mature animals (∼6 mon old at start of irradiation)(23, 26–37). Second, type of behavioral test matters. Recent work with mature rodents shows HZE particle exposure decreases performance in some - but not all - behavioral tests, and even tests that engage similar neural circuits produce distinct results (23, 26–31, 38). A potential contribution to these test-dependent discrepancies is the testing environments used for each task. In humans (and astronauts), automated computerized cognitive assays help control for the influence of testing environments (39–43), but such an approach has not been used to assess cognitive performance in rodents after HZE exposure. Third, breadth of testing matters. Preclinical studies on space radiation typically assess one or two cognitive domains (24, 44–47). In contrast, astronauts repeatedly undergo test batteries - often on a touchscreen platform - to assess integrity of many cognitive domains over time (39, 48, 49). To this end, many aspects of neuroscience have employed rodent touchscreen testing, a platform extensively validated for its ability to provide multidimensional assessment of functional integrity of brain circuits in a highly-sensitive and translationally-relevant way (50–61). Given the power of touchscreen testing, it is surprising that the effect of space radiation on a battery of rodent touchscreen tests is unknown. This is particularly notable as the touchscreen platform permits analysis of many higher cognitive functions - such as pattern separation - which are part of the astronaut’s mission-critical skill set yet which have not been preclinically assessed for their sensitivity to space radiation.

To address these major knowledge gaps, mature C57BL/6J male mice received either Sham irradiation (IRR) or whole body ^56^Fe particle IRR and were assessed on a battery of touchscreen cognitive tasks (54, 56, 62–65). Sham and ^56^Fe IRR mice performed similarly in touchscreen tasks of complex learning, cognitive flexibility, visuospatial learning, and stimulus-response habit learning. Notably, ^56^Fe IRR mice performed better than Sham in location discrimination, a touchscreen task of pattern separation ability, as they discriminated similar visual cues in fewer days and more accurately than Sham mice. This improvment was not restricted to ^56^Fe IRR or to appetitive testing; mice exposed to either ^56^Fe or ^28^Si also discriminated contexts faster and more consistently relative to Sham mice when assessed on a classical, non-touchscreen task of pattern separation: fear-based contextual discrimination fear conditioning (CDFC). These data show whole body exposure to HZE particles is not detrimental to high level cognition in mice and actually enhances performance in certain mission-critical tasks, such as pattern separation.

## RESULTS

### Mice exposed to whole body ^56^Fe radiation demonstrate overall normal perceptual discrimination, association learning, and cognitive flexibility in touchscreen testing

Whole body ^56^Fe IRR was delivered via fractionation (Frac; 3 exposures of 6.7 cGy every other day, total 20 cGy) to male C57BL/6J mice at 6 mon of age. This total dose is submaximal to that predicted for a Mars mission (9, 66), and the fractionation interval (48 hours [hr]) was based on the importance of the inter-fraction period for potential repair processes (67–69). As previously reported (70–72), this dose and these fractionation parameters do not interfere with weight gain or cause hair loss (Fig. S1A).

Beginning 1 mon post-IRR, Sham and ^56^Fe IRR mice began training on a touchscreen platform extensively validated in rodents (54, 56, 64, 73–75)(Fig. 1A). Mice initially went through five stages of general touchscreen training (Fig. 2A), with performance reflecting instrumental or operant learning. Sham and ^56^Fe IRR mice completed most stages of the initial operant touchscreen training in similar periods of time (Fig. 2A). The exception was the final stage, Punish Incorrect (PI, an incorrect trial results in a timeout period); on average, ^56^Fe IRR mice finished PI in ∼40% fewer days relative to Sham mice (Fig. 2A, Table S1).

**Figure 1.**
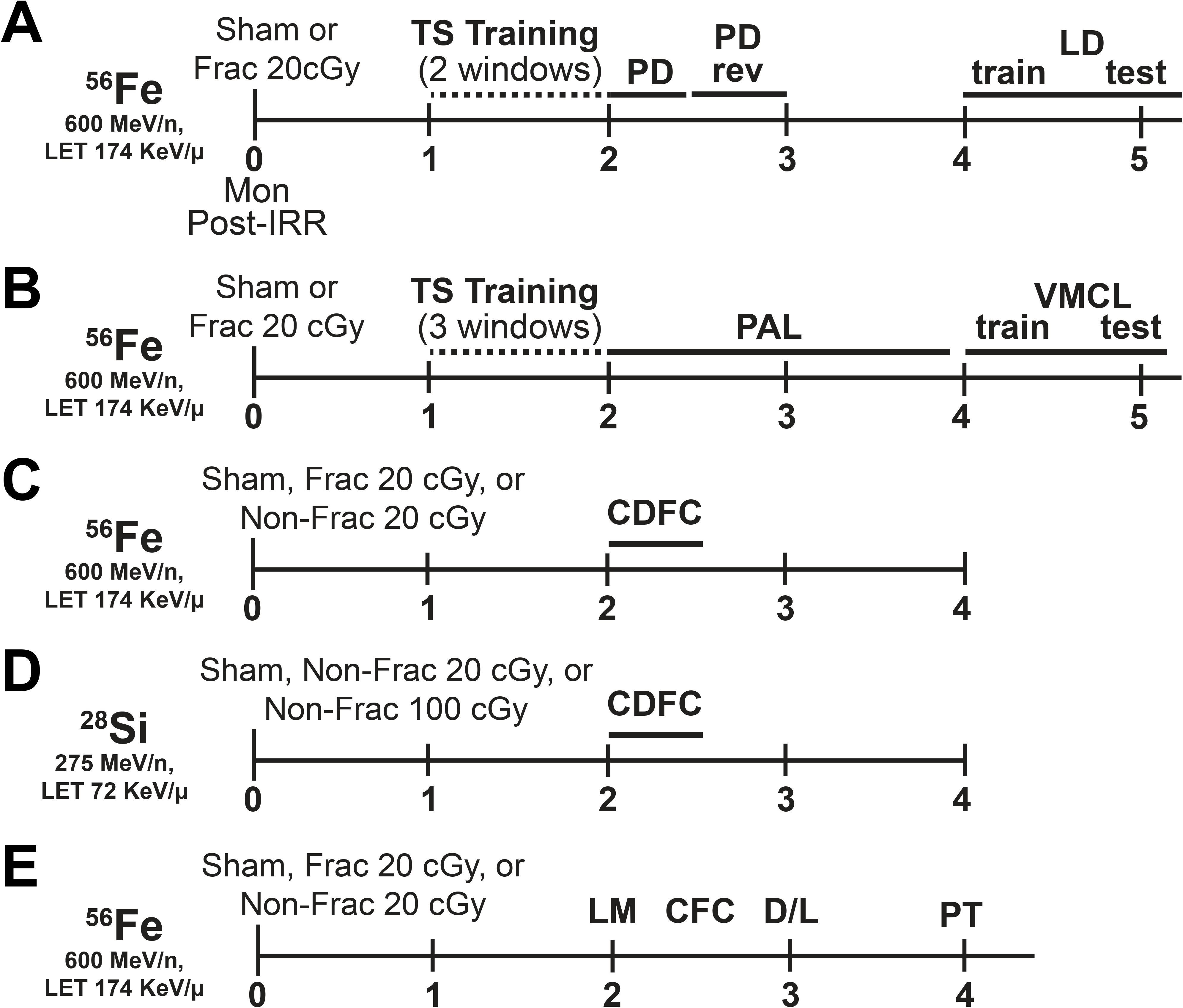
Timeline of experimental groups and behavior tests. **(A-E)** C57BL/6J male mice (JAX cat#00684) received whole-body exposure to particles of ^56^Fe **(A-C, E)**, ^28^Si **(D)**, or Sham exposure at 6-months (mon) of age (0-mon post-irradiation (IRR)). Mice subsequently were run on a variety of touchscreen behavioral tests (**A:** TS training, PD, PD rev, LD, **B:** TS training, PAL and VMCL) or classic behavior tests (**C-D:** CDFC, **E:** LM, CFC, D/L, PT) and DCX^+^ cells were quantified 4-mon post-IRR **(C)**. CDFC=contextual discrimination fear conditioning, CFC=contextual fear conditioning, D/L=dark/light box test, IRR=irradiation, LD=location discrimination, LM=locomotor, mon=months, PAL=paired associates learning, PD=pairwise discrimination, PD rev=PD reversal, PT=pain threshold, VMCL=visuomotor conditioning learning, TS=touchscreen.

**Figure 2.**
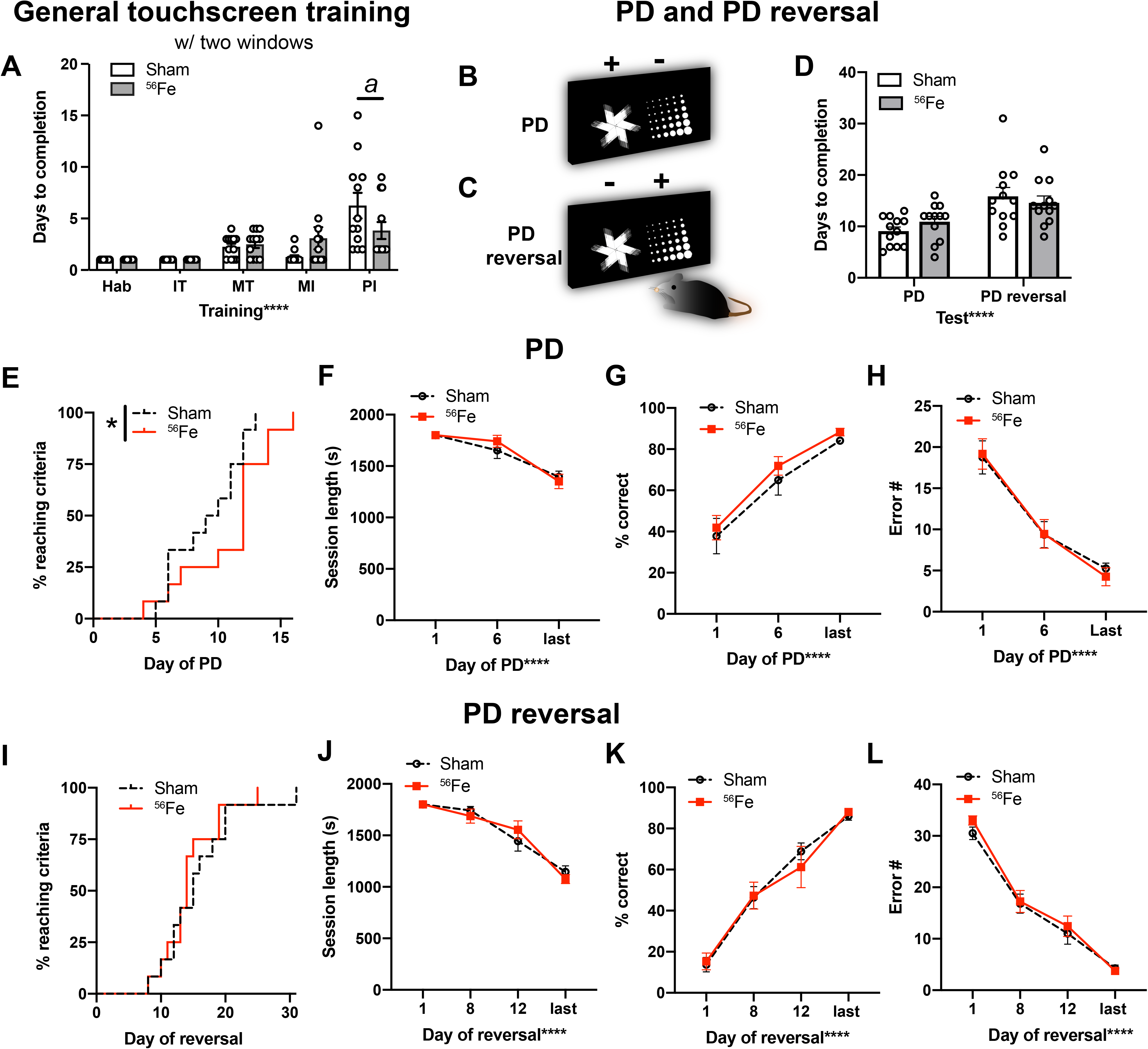
Mice exposed to whole body ^56^Fe IRR at 6-month of age complete the final stage of general touchscreen training in fewer days compared to Sham mice, but perform similarly to Sham mice overall in the Pairwise Discrimination (PD) and reversal (PD rev). **(A)** Sham and ^56^Fe IRR mice performed similarly in the first four steps of general touchscreen training with two windows: Habituation (Hab), Initiate Touch (IT), Must Touch (MT), and Must Initiate (MI). However, ^56^Fe IRR mice completed the Punish Incorrect (PI) stage of general touchscreen training in fewer days than Sham mice. **(B-C)** Sample touchscreen images for PD and PD reversal tests. **(D)** Sham and ^56^Fe IRR mice completed PD and PD rev in similar number of days. **(E)** Cumulative distribution function showing the difference in the rate of days required to complete PD between Sham and ^56^Fe IRR mice. **(F-H)** Sham and ^56^Fe IRR mice performed similarly in PD (**F**: session length, **G**: percent(%) correct, **H**: Error number(#)). **(I)** Cumulative distribution function showing no difference in the test days required to complete PD rev between two groups. **(J-L)** Sham and ^56^Fe IRR mice performed similarly in PD rev (**J:** session length, **K:** % correct, **L:** Error #). Mean±SEM. Two-way RM ANOVA in **A, D, F-H, J-L**, *p<0.05, ****p<0.0001, post hoc: Bonferroni *a* p<0.05 in Sham vs. ^56^Fe; Mantel-Cox test **(E,I)**, *p<0.05. s=seconds.

Mice then advanced to pairwise discrimination (PD, visual discrimination) and PD reversal (reversal learning, Fig. 2B, 2C), tests which reflect perceptual discrimination and association learning as well as cognitive flexibility, respectively, and rely on cortical (prefrontal, orbital frontal, perirhinal) and striatal circuits (56, 73, 74). On average, both ^56^Fe IRR and Sham mice completed PD and PD reversal in a similar number of days (Fig. 2D, Table S1). However, analysis of the distribution of subjects to reach criteria each day revealed significant difference between Sham and ^56^Fe IRR mice (Fig. 2E). Specifically, 50% of Sham mice reached PD completion criteria at 9.5 days, while 50% ^56^Fe IRR mice reached criteria at 12 days. However, Sham and ^56^Fe IRR mice did not differ with regard to average session length, percent correct, or number of errors (Fig. 2F-H). In PD reversal, the distribution of subjects to reach completion criteria was not different between Sham and ^56^Fe IRR mice (Fig. 2I), with 50% of Sham and ^56^Fe IRR mice reaching PD reversal completion criteria at 15 and 14 days, respectively (Fig. 2I). As with PD, Sham and ^56^Fe IRR mice did not differ in regard to PD reversal average session length, percent correct, or number of errors (Fig. 2J-L).

### Mice exposed to whole body ^56^Fe demonstrate normal visuospatial learning and stimulus-response habit learning in touchscreen testing

A parallel group of mice was used to assess the influence of ^56^Fe IRR object-location paired associates learning (PAL) and visuomotor conditional learning (VMCL) which reflect visuospatial and stimulus-response habit learning, respectively, and rely on intact circuits of the hippocampus (56, 63, 64)(PAL) and striatum and posterior cingulate cortex (56, 64, 75)(VMCL)(Fig. 1B). Consistent with results in the first cohort of mice, Sham and ^56^Fe IRR mice completed most stages of the initial operant touchscreen training in similar periods of time (Fig. 3A), again with the exception of PI where ^56^Fe IRR mice finished in ∼20% fewer days than Sham.

**Figure 3.**
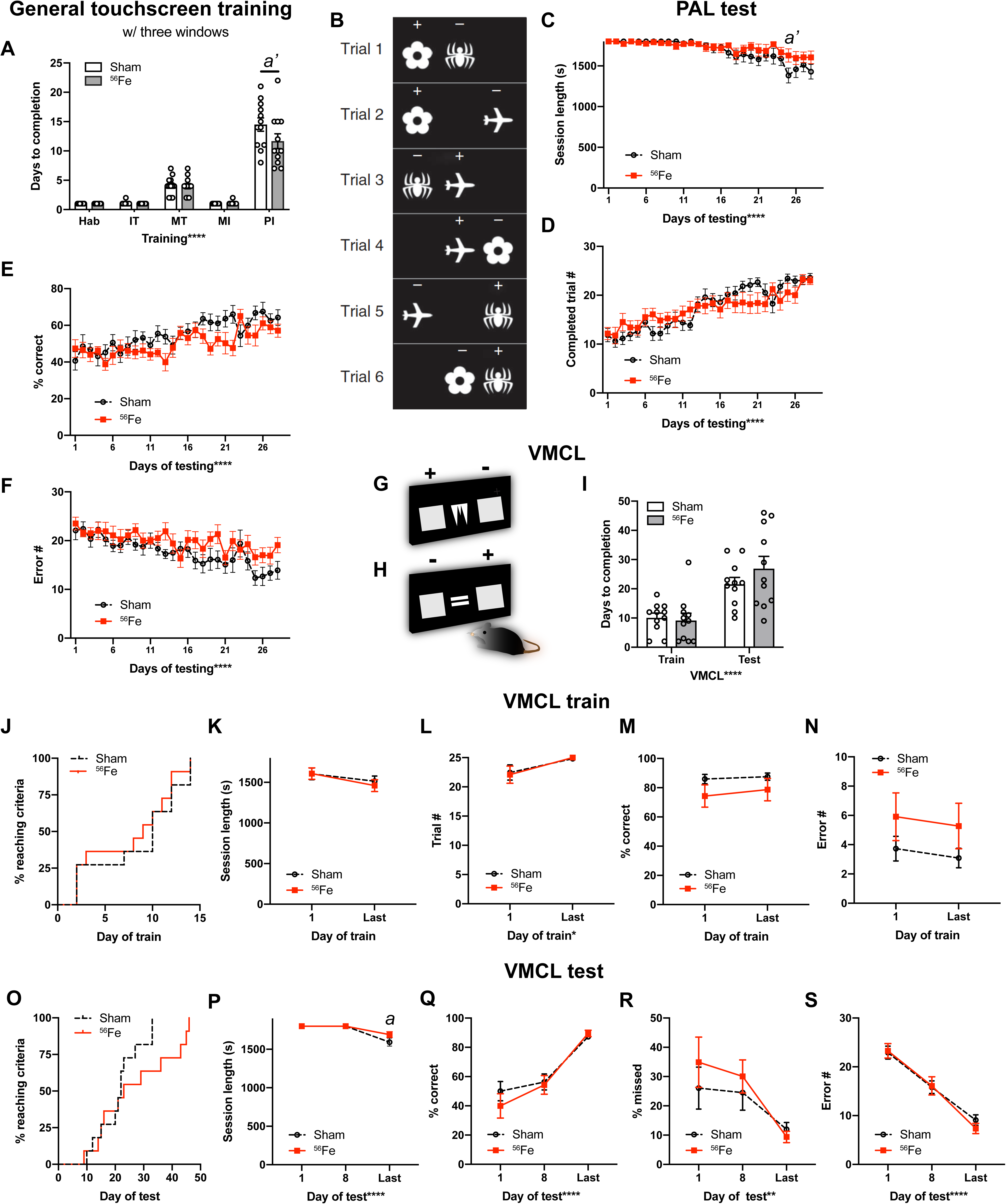
Mice exposed to whole body ^56^Fe IRR at 6-month of age complete the final stage of general touchscreen testing in fewer days than Sham mice, but perform similarly to Sham in tests of rule-based learning and stimulus-response habit learning. **(A)** Sham and ^56^Fe IRR mice performed similarly in the 4 first steps of general touchscreen training stages with three windows, including Habituation (Hab), Initiate Touch (IT), Must Touch (MT), and Must Initiate (MI). However, ^56^Fe IRR mice completed the Punish Incorrect (PI) stage of general touchscreen training in fewer days than Sham mice. **(B)** Sample touchscreen images for different Paired Associates Learning (PAL). **(C-F)** Sham and ^56^Fe IRR mice performed similarly in PAL (**C**: session length, **D**: completed trials, **E**: percent (%) correct, **F**: Error number (#)). **(G-H)** Sample touchscreen images for Visuomotor Conditional Learning (VMCL) train and test phases. **(I)** Sham and ^56^Fe IRR mice performed similarly in VMCL train and test. **(J)** Cumulative distribution function showed no difference in days required to complete training. Distribution of Sham and ^56^Fe IRR mice (n=11/group) did not differ in days required to complete VMCL training. **(K-N)** Sham and ^56^Fe IRR mice performed similarly in VMCL train (**K**: session length, **L:** completed trials, **M:** % correct, **N:** Error #). **(O)** Cumulative distribution function showed no difference in days required to complete VMCL test. Distribution of Sham and ^56^Fe IRR mice (n=11/group) did not differ in VMCL test. **(P-S)** Sham and ^56^Fe IRR mice performed similarly in VMCL test (**P**: session length, **Q:** % correct, **R:** % missed, **S:** Error #). Mean±SEM. Two-way RM ANOVA in **A, C-F, I, K-N, P-S**, *p<0.05, **p<0.001, ****p<0.0001, post hoc: Bonferroni a p<0.05, a’ p<0.01 in Sham vs. ^56^Fe; Mantel-Cox test in **J,O.** PAL=different Paired Associates Learning, s=seconds, VMCL=Visuomotor Conditional Learning.

In PAL (Fig. 3B), Sham and ^56^Fe IRR mice were similar in session length, number of trials, percent correct, and number of errors over the 29-day testing period (Fig. 3C-F). In both VMCL train and test (Fig. 3G, 3H), Sham and ^56^Fe IRR mice had similar average days to completion (Fig. 3I). In VMCL train, Sham and ^56^Fe IRR mice performed similarly in regard to distribution of subjects to reach criteria each training day (50% subjects reached criteria at 10 days in Sham mice vs. 9 days in ^56^Fe IRR mice), session length, number of trials, percent correct, and number of errors (VMCL train; Fig. 3J-N, Table S1). In VMCL test, Sham and ^56^Fe IRR mice had similar distribution of subjects to reach criteria each training day (50% of subjects reached criteria at 22 days in Sham mice vs. 23 days in ^56^Fe IRR mice), session length, number of trials, percent correct, and number of errors (VMCL test; Fig. 3O, Q-S, Table S1). However, the time to complete the session on the last day of VMCL test was longer in ^56^Fe IRR relative to Sham mice (Fig. 3P).

### Whole body ^56^Fe IRR exposure improves pattern separation in an appetitive-based location discrimination touchscreen task

The brain region most studied with regard to space radiation-induced deficits in function and activity-dependent processes (i.e. neurogenesis) is the hippocampal dentate gyrus (70–72, 76–78). Therefore, we hypothesized whole body ^56^Fe IRR impairs pattern separation, a cognitive function reliant on dentate gyrus integrity (79–81). To determine the effect of HZE radiation on pattern separation, Sham and ^56^Fe IRR mice were assessed on a touchscreen location discrimination (LD) task (54)(Fig. 1A). In the LD training portion of the assessment (LD train, Fig. 4A), Sham and ^56^Fe IRR mice had similar distribution of the proportion of subjects reaching criteria (Fig. 4B), average days to completion, session length, and percent correct (Fig. 4C-E). However, Sham and ^56^Fe IRR mice differed in LD performance (LD test, Fig. 4F) in several aspects. First, the distribution of proportion of subjects reaching criteria was distinct in ^56^Fe IRR mice vs. Sham mice (Fig. 4G). ^56^Fe IRR mice reached criteria at >3x faster rate vs. Sham mice, and 50% of ^56^Fe IRR mice reached criteria by 4 days vs. Sham mice reaching criteria by 6 days. Second, ^56^Fe IRR mice completed LD test in fewer days than Sham mice (Fig. 4H), although both groups showed similar session length and number of completed trials (Fig. 4I-J). Third, ^56^Fe IRR mice performed LD test more accurately than Sham mice both overall (Fig. 4K) as well when presented with stimuli separated by either large or small distances (Fig. 4L).

**Figure 4.**
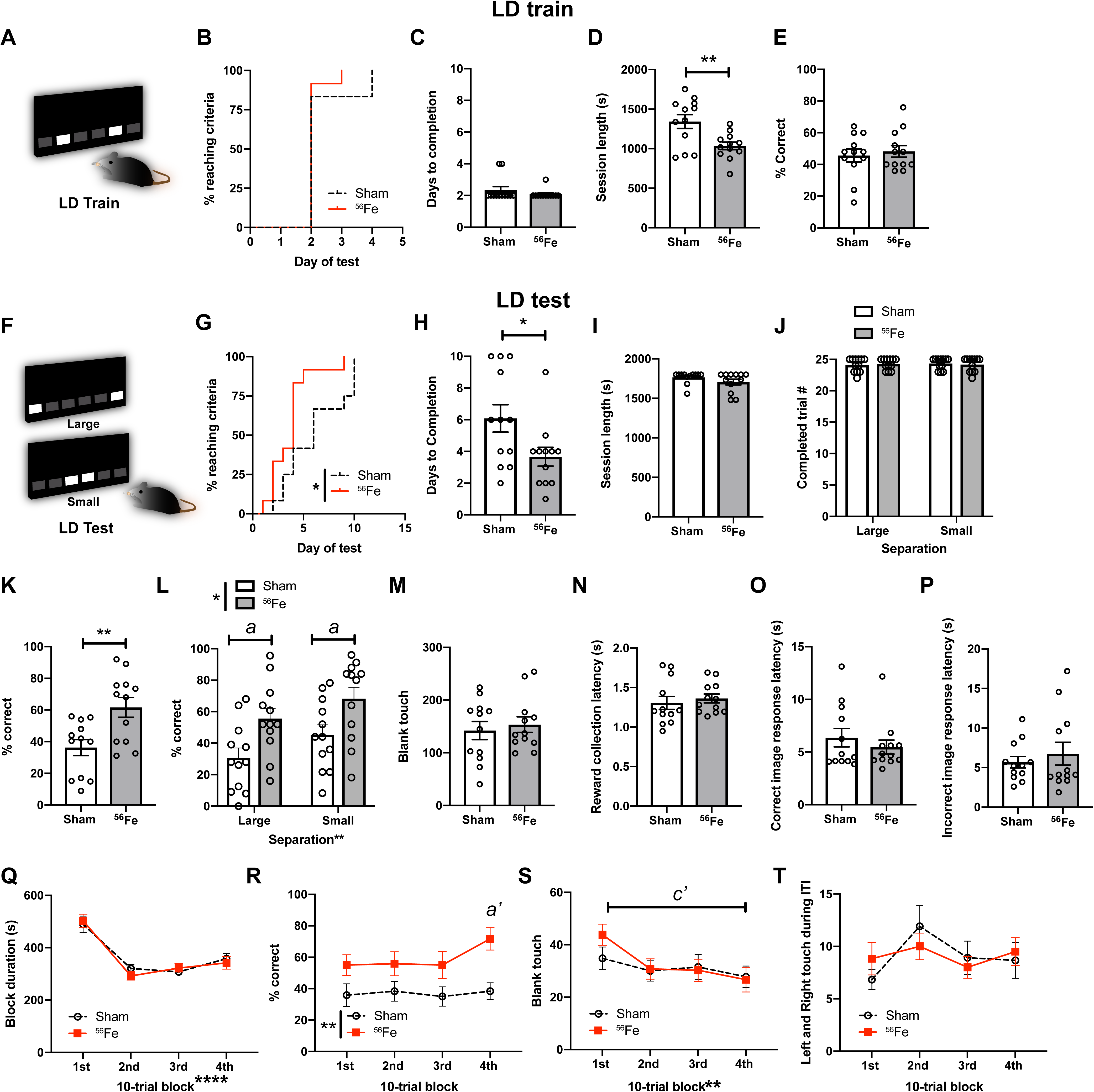
On an appetitive pattern separation task, mice exposed to whole body ^56^Fe IRR at 6-month of age distinguish two similar visual cues earlier and with greater accuracy on the last test day relative to Sham mice. **(A)** Sample touchscreen images for location discrimination training (LD train). **(B-E)** Sham and ^56^Fe IRR mice performed similarly in LD train. **B:** distribution of subjects reaching criteria, **C:** days to completion, **D:** session completion time, **E:** % correct. **(F)** Sample touchscreen images for LD testing (LD test). **(G-J)** ^56^Fe IRR mice completed the LD test earlier than Sham **(G,H)**, but no difference in session completion time **(I)** or number of completed trials **(J)**. **(K-L)** ^56^Fe IRR mice were more accurate overall **(K)** and on both “Large” and “Small” separation trials compared to Sham mice **(L)**. **(M-P)** Sham and ^56^Fe IRR mice made similar number of blank touches to non-stimuli windows **(M)** and had similar reward collection latency **(N)**, correct image response latency **(O)**, and incorrect image response latency **(P)**. **(Q-T)** Sham and ^56^Fe IRR mice had similar block duration in each 10-trial block **(Q)**. However, ^56^Fe IRR mice had higher accuracy in the 4th 10-trial block (31st −40th trial) compared to Sham mice **(R)**. Sham and ^56^Fe IRR mice made similar number of blank touches in each block **(S)** and left and right touches during inter-trial interval (ITI) **(T)**. Mean±SEM. Mantel-Cox test, *p<0.05 in **B, G**; Unpaired, two-tailed t-test in **C-E, H-I, K, M-P**; Two-way RM ANOVA,*p<0.05, **p<0.01, post hoc: Bonferroni in **J, L, Q-T**, *a* p<0.05, *a’* p<0.01 in Sham vs.^56^Fe mice in **L, R**, c’ p<0.01 1st and 4th block in ^56^Fe mice in **S**. s=seconds.

We next behaviorally probed reasons why ^56^Fe IRR mice had improved pattern separation relative to Sham mice. For example, the improved location discrimination in ^6^Fe IRR mice may be reflective of unintentional screen touches, perhaps due to IRR-induced alteration of attention to stimuli or motivation to obtain reward. However, the number of blank touches (Fig. 4M), reward collection latency (Fig. 4N), and choice latency (Fig. 4O, 4P) were not different between ^6^Fe IRR mice and Sham mice. Also, since the location of the rewarded stimuli changed daily but maintained within each session, it is possible that pattern separation is progressively improved within a session, particularly on the last test day. Sham and ^56^Fe IRR mice had similar last day block duration and left/right touches during intertrial interval, but ^56^Fe IRR mice had a greater percent correct during the 4th 10-trial block relative to Sham mice (Fig. 4Q, 4R, 4T). In addition, while Sham mice did not differ between the 1st and 4th 10-trial blocks on the last day, ^56^Fe IRR mice had fewer blank touches in the 4th 10-trial block relative to the 1st 10-trial block. These data suggest that on the last day of LD, ^56^Fe IRR mice demonstrate within-session enhanced pattern separation (Fig. 4S).

### Whole body ^56^Fe and ^28^Si IRR exposure improves pattern separation in a foot-shock based contextual discrimination task

To assess whether ^56^Fe IRR-induced improvement in pattern separation was restricted to appetitive tasks, a parallel cohort of mice was exposed to Sham or ^56^Fe IRR and tested on pattern separation using a classic pattern separation behavior paradigm: contextual discrimination fear conditioning (CDFC)(80, 82–85). To specifically assess whether particle delivery influenced behavioral outcome, 6-mon-old C57BL/6J mice received either Sham IRR, whole body ^56^Fe IRR via fractionation (Frac 20 cGy; 3 exposures of 6.7 cGy), or whole body ^56^Fe IRR via non-fractionation (Non-Frac 20 cGy; 1 exposure of 20 cGy; Fig. 1C). As previously reported (70–72), Sham IRR, Frac 20 cGy, and Non-Frac 20 cGy mice had similar weight changes over time (Fig. S1A).

Beginning ∼2-mon post-IRR (8 mon of age), mice underwent CDFC (Fig. 5, Fig. S2) to learn that one context (Context A) was paired with a foot shock while another similar context (Context B) was a non-shock context. When tested in CDFC, Sham mice discriminated the two contexts by Days 9-10 (Block 5), as they froze more in the shock-paired context (Context A) compared to the non-shock context (Context B; Fig. 5A, Table S1). However, mice exposed to either Frac 20 cGy or Non-Frac 20 cGy of ^56^Fe IRR discriminated the contexts by Days 3-4 (Block 2, Fig. 5B, 5C, Table S1). Direct comparison across treatment groups revealed Frac 20 cGy and Non-Frac 20 cGy mice froze more in Context A vs. Context B in Blocks 2 and 4, earlier than Sham (Fig. 5D-F, Table S1). Possible explanations for these results include differential activity, anxiety, or pain sensitivity in Sham vs. ^56^Fe IRR mice. To address these possibilities, parallel groups of mice underwent assessment for locomotion (Fig. S1B), dark/light testing (Fig. S1C, S1D) and pain threshold (Fig. S1E-G). However, Sham, Frac, and Non-Frac mice performed similarly on all these tests (Fig. S1B-G). Thus, both Frac and Non-Frac 20 cGy ^56^Fe IRR mice learned to pattern separate earlier relative to Sham mice without overt changes in locomotion, anxiety-like behavior, or sensitivity to pain.

**Figure 5.**
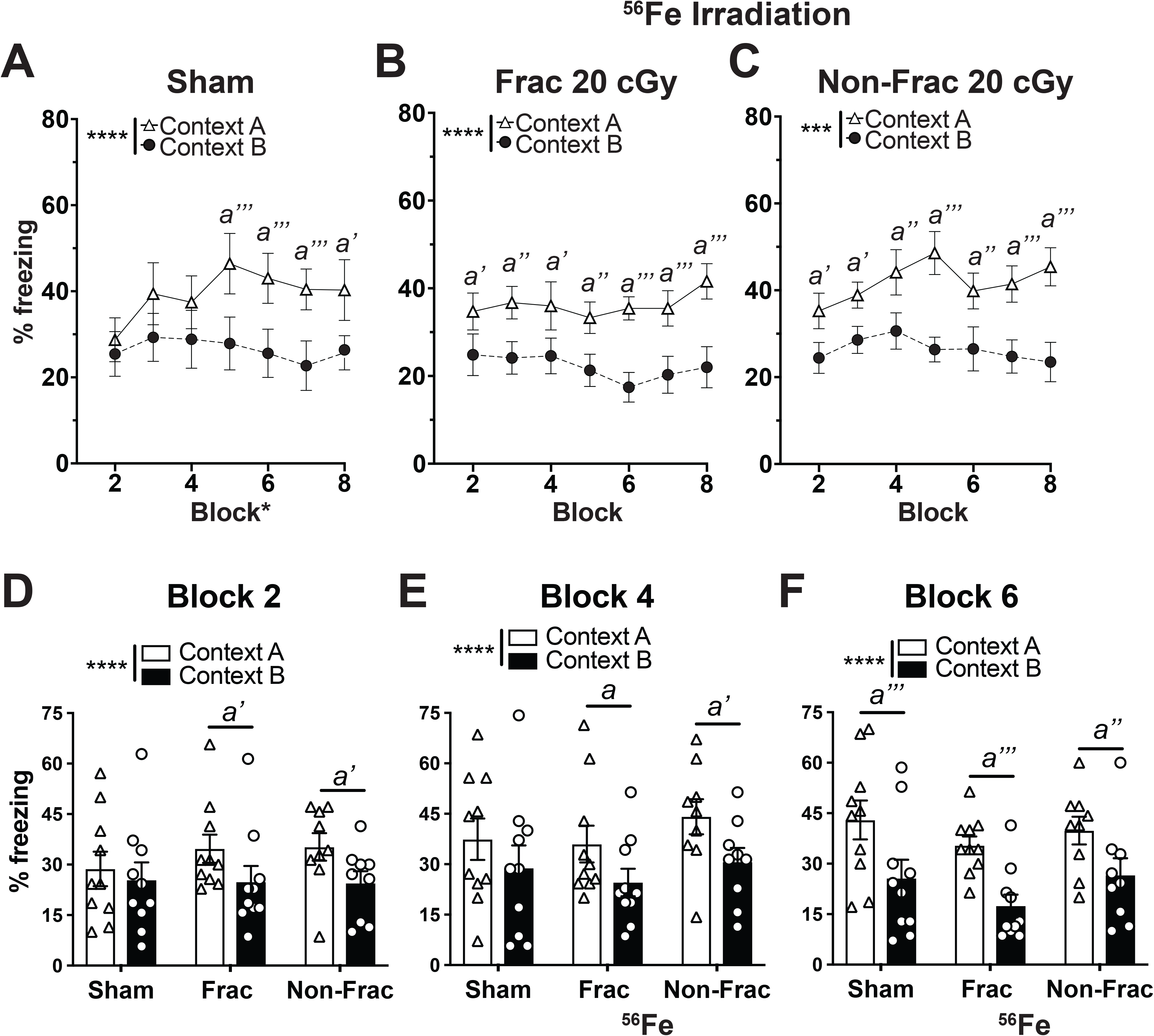
On an aversive pattern separation test, mice exposed to whole body ^56^Fe IRR at 6-month of age discriminate two contexts earlier than mice exposed to Sham IRR. **(A)** Sham mice discriminate Context A (shock context) from Context B (non-shock context) by Block 5. **(B-C)** Frac **(B)** and Non-Frac **(C)** ^56^Fe mice discriminate Context A from Context B by block 2. **(D-F)** When examined at Block 2 **(D)**, Block 4 **(E)**, and Block 6 **(F)**, Frac and Non-Frac ^56^Fe discriminate by Block 2. Mean±SEM. **(A-F)** Two-way RM ANOVA, *p<0.05, **p< 0.01, ***p< 0.001, ****p< 0.0001, Bonferroni post-hoc tests in **A-F**. *a* p<0.05, *a’* p<0.01, *a’’* p<0.001, *a’’’* p<0.0001 in Context A vs B.

To determine if the improvement in CDFC pattern separation generalized to other fear-based hippocampal- and amygdala-based learning, a parallel cohort of mice received Sham or ^56^Fe IRR and underwent classical contextual fear conditioning (CFC; Fig. 1E, Fig. S3A, S3B). Sham and ^56^Fe IRR mice (both Frac and Non-Frac 20 cGy groups) performed similarly in the context test (Fig. S3C) and in the cue test both pre-tone and during tone (Fig. S3D). Importantly, to see if the space radiation-induced improvement in CDFC was dependent on the type of heavy particle used, CDFC was also performed with mice exposed to whole body ^28^Si IRR (Fig. 1D, 6), a particle with a smaller track structure than ^56^Fe (86). Sham mice spent more time freezing in Context A vs. Context B only on Days 9-10 (Block 5) and Days 15-16 (Block 8) (Fig. 6A, D-F). Mice exposed to 20 cGy of ^28^Si discriminated between the two contexts as early as Days 11-12 (Block 6; Fig. 6B, D-F). Notably, mice exposed to 100 cGy of ^28^Si were able to discriminate between the two contexts as early as Days 5-6 (Block 3; Fig. 6C, D-F). Taken together, these data show that exposure to two different HZE particles - either ^56^Fe or ^28^Si - results in earlier separation ability relative to Sham mice on the shock-based CDFC pattern separation test.

**Figure 6.**
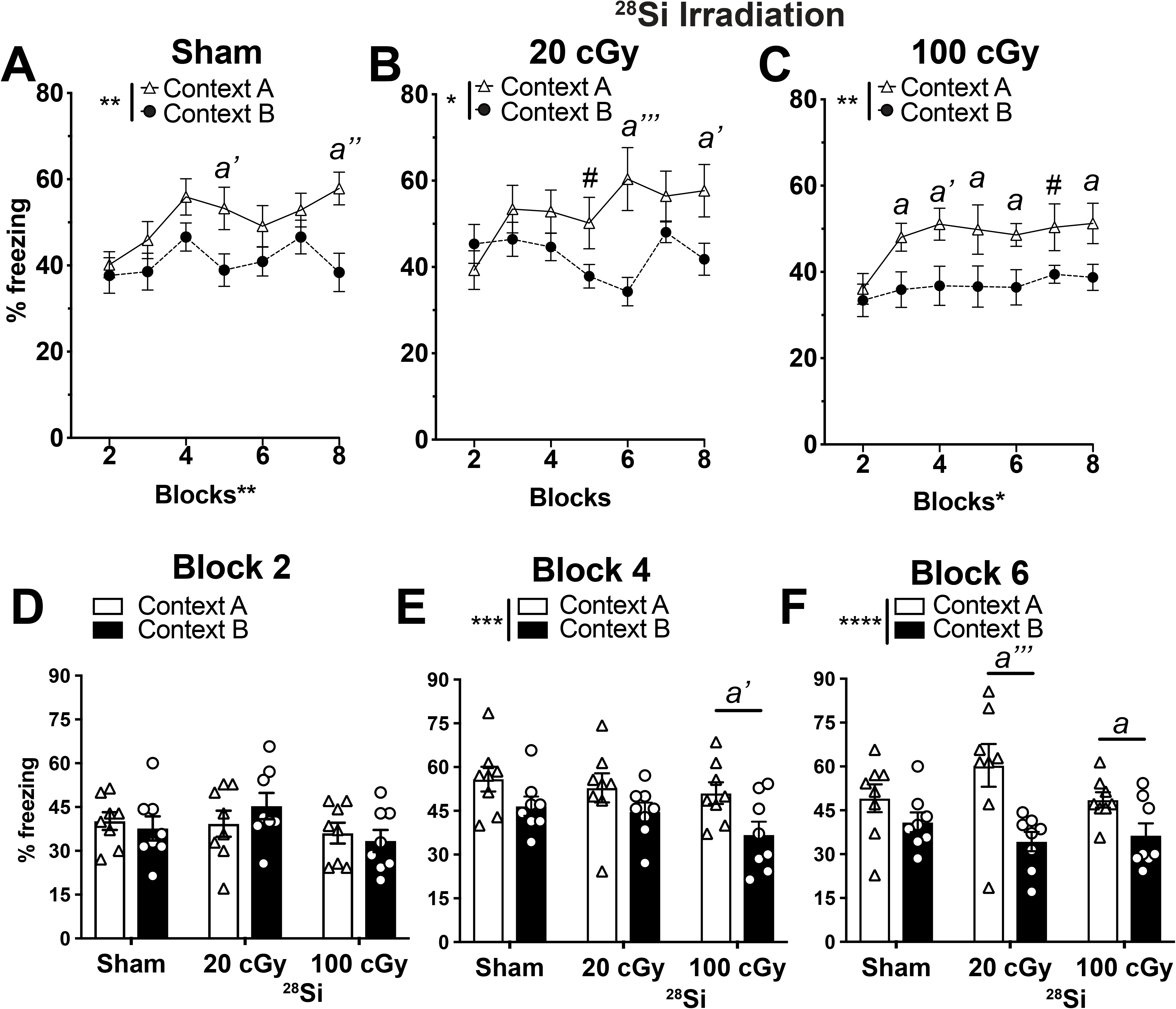
On an aversive pattern separation test, mice exposed to a different HZE particle - ^28^Si- at 6 month of age also discriminate two contexts earlier than mice exposed to Sham IRR. **(A)** Sham mice discriminate Context A (shock context) from Context B (non-shock context) by Block 5. **(B-C)** While 20cGy ^28^Si mice **(B)** discriminate Context A from Context B by Block 5, 100cGy ^28^Si mice **(C)** discriminate by Block 3. **(D-F)** When examined at Block 2 **(D)**, Block 4 **(E)**, and Block 6 **(F)**, 100cGy Si mice by Block 4 and both 20cGy and 100cGy ^28^Si mice discriminate by Block 6. Mean±SEM. **(A-F)** Two-way RM ANOVA, *p<0.05, **p< 0.01, ***p< 0.001, ****p< 0.0001, Bonferroni post-hoc tests in **A, B, C, E, F**, #0.06<p<0.05, *a* p<0.05, *a’* p<0.01, *a’’* p<0.001, *a’’’* p<0.0001 in Context A vs. B.

### 56Fe IRR decreases neurogenesis 4 mon post-IRR

Pattern separation ability is dependent on new dentate gyrus neurons as well as dentate gyrus activity, and an inducible increase in adult neurogenesis improves pattern separation (80, 87, 88). To assess whether the IRR-induced improvement in pattern separation reported here was correlated with increased neurogenesis, we used stereology to quantify the number of cells in the dentate gyrus immunoreactive for doublecortin (DCX, Fig. 7A), a widely-accepted marker for neurogenesis (89–91). Although mice exposed to either Fractionated or Non-Fractionated ^56^Fe radiation had improved context discrimination compared to control mice (Fig. 5), these mice had fewer DCX+ cells compared to control mice (Fig. 7B-C, Table S1).

**Figure 7.**
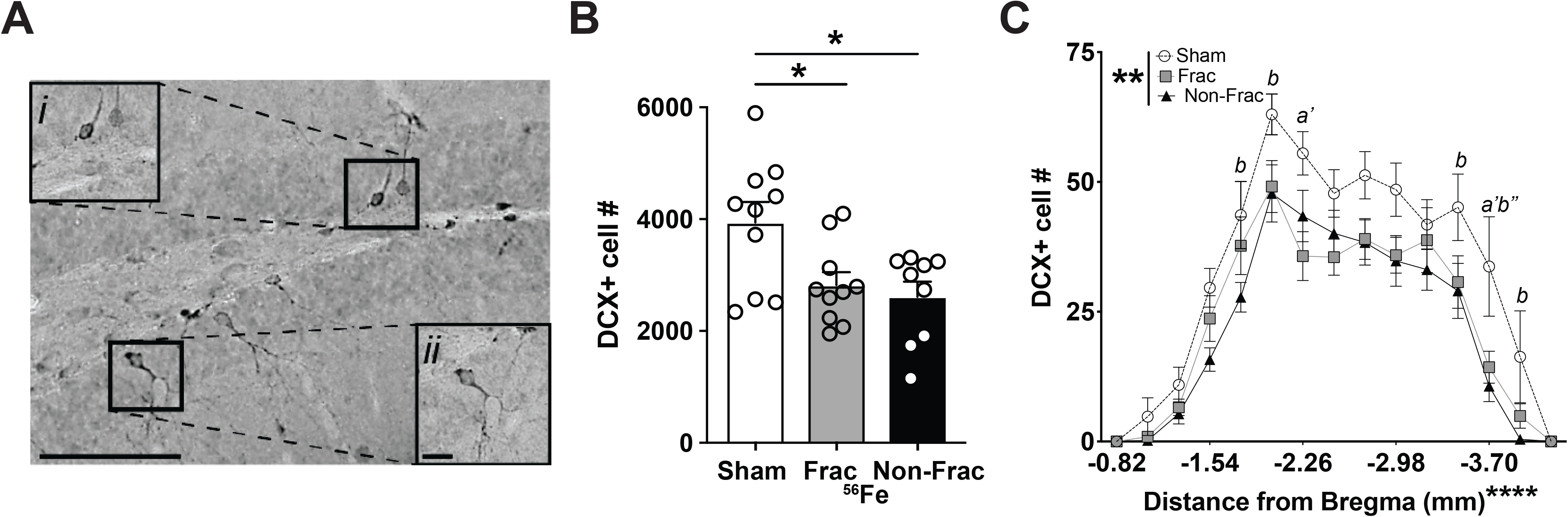
Stereological quantification reveals fewer immature dentate gyrus neurons (doublecortin (DCX)+ cells) 4 months post-whole body ^56^Fe particle radiation relative to Sham mice. **(A)** Representative photomicrograph of DCX+ cell in the mouse dentate gyrus subgranular zone. Insets: higher magnification of boxed areas in main image. Scale bar=100 um in **A**, 10 um in inset *ii*. **(B-C)** Relative to Sham mice, Frac, and Non-Frac ^56^Fe mice have fewer DCX+ dentate gyrus cells. Mean±SEM. **(B)** One-way ANOVA Bonferroni posthoc. *p<0.05, **(C)** Two-way ANOVA, Bonferroni posthoc. *a’* p<0.01 Sham vs. Frac, *b* p<0.05, *b”* p<0.001 Sham vs. Frac. Frac=fractionation, Non-Frac=non-fraction.

## DISCUSSION

Astronaut training and in-mission assessment rely on touchscreen testing due to its flexibility in probing a variety of cognitive functions. Rodent touchscreen testing similarly allows researchers to probe the multidimensional functional integrity of brain circuits in a highly-sensitive and translationally-relevant way (50–56), but prior to the present work it was unknown how space radiation influences touchscreen performance. Based on the large literature with young animals (92–94) and the negative impact of HZE particle exposure on the central nervous system (22, 95), we hypothesized whole-body exposure to ground-based HZE particles would diminish the performance of mice in touchscreen-based behaviors, particularly those behaviors reliant on the dentate gyrus, such as pattern separation. The results of our multi-domain cognitive assessment showed our hypothesis was wrong. Mature mice exposed to either Sham IRR or HZE particles performed similarly in touchscreen tasks of visual discrimination, cognitive flexibility, rule-based learning, and object-spatial associated learning, in classical hippocampal- and amygdala-based tasks (i.e. CFC), and in tasks that detect anxiety-like behavior (i.e. D/L). Surprisingly, IRR mice performed better than Sham IRR mice in pattern separation tasks when assessed on either appetitive (LD test) or aversive (CDFC) platforms. Thus, our study suggests whole body exposure of HZE particles in maturity is not detrimental to high-level cognition, and actually enhances performance in the mission-critical task of pattern separation.

There are three aspects of the present results that are notable from the perspective of behavioral neuroscience in general, and multiple memory systems in particular (96, 97). First, in both humans and rodents, hippocampal damage can actually facilitate behavioral performance on certain tasks (98–100). For example, when amnesic patients with partial hippocampal injury are given extended exposure to study materials, they can improve their recognition memory to the level of control subjects. Such an improvement is not seen after severe hippocampal injury. Thus, it is reasonable to consider whether the improved pattern separation ability presented here result from HZE particle-induced partial damage to the hippocampus. This is unlikely, as the HZE particle parameters used here do not induce detectable damage to post-mitotic neurons in the adult rodent brain (101–105) or, as shown here, deficits in other tasks that engage the hippocampus (PAL, CFC).

Second, as memory mechanisms in the medial temporal lobe (i.e. hippocampus) and basal ganglia (i.e. dorsal striatum) may sometimes compete (97), it is possible the improved dentate gyrus-based pattern separation reported here is associated with decreased dorsal striatum-based ‘habit’ learning. However, we find pattern separation is improved in ^56^Fe relative to Sham mice without a change in VMCL habit learning, suggesting normal dorsal striatal function. Finally, the improved pattern separation reported here is reminiscent of the excessive attention seen in some psychiatric disorders - such as autism or obsessive compulsive disorder (OCD) - and in animal models for these disorders (63, 106–108). Evaluation of autistic- or OCD-like behavioral patterns after HZE particle exposure using other touchscreen paradigms (i.e. extinction, 5-choice serial reaction time test, 5-choice continuous performance reaction task) would clarify whether the improved pattern separation ability demonstrated here is accompanied by maladaptive behaviors (i.e. impaired attention and increased impulsivity)(109–111).

What might be the neural mechanism underlying the improved pattern separation in HZE-irradiated mice reported here? One possibility is an HZE-induced shift in underlying brain circuit activity. In rodents and humans, pattern separation requires the appropriate balance of activity in the entorhinal cortex-dentate gyrus network (79, 87, 112–115). In aged humans, a decline in pattern separation (116–118) is proposed to be due to a hypoactive anterolateral enthorhinal cortex and hyperactive dentate gyrus/CA3 (118). Thus, it is possible the HZE-induced improved pattern separation reported here in mouse results from an opposite activity shift: a hyperactive enthorhinal cortex and hypoactive dentate gyrus/CA3. Indeed, in rodents, pattern separation performance is correlated with dentate gyrus activity; better performance results in a hypoactive dentate gyrus, and worse performance results in a hyperactive dentate gyrus (80, 112). As pattern separation engages distinct hippocampal networks relative to other hippocampal-dependent tests (such as novel object recognition)(119–121), such an HZE-induced shift in hippocampal networks may explain why we see improved pattern separation - while other groups see decreased novel object recognition - after HZE exposure.

Another possibility is that the improved pattern separation we report in HZE-irradiated mice is due to HZE-induced conditions in the dentate gyrus that favor “sparse encoding” of entorhinal cortical input. Sparse encoding in dentate gyrus granule cell neurons is critical for pattern separation, as it minimizes interference between memory representations of similar but not identical experiences (122–126). This sparsity is due in part to inhibition of dentate gyrus granule cell neurons by GABAergic interneurons and mossy cells (127–130). It is unknown how the HZE particle parameters used here influence dentate gyrus GABAergic interneurons and mossy cells in mature mice. However, exposure to other energetic particles that comprise space radiation alters the inhibitory network in the dentate gyrus and other hippocampal subregions of young adult rodents (131, 132). In the future, evaluation of GABAergic signaling and other measures relevant to sparse encoding (e.g. number and functionality of hilar interneurons and mossy cells, pattern of memory-induced immediate early gene activation) (133) after Mars-relevant exposure to space radiation would allow testing of the hypothesis that HZE-induced improvement in sparse encoding contributes to the HZE-induced improvement in pattern separation reported here.

A third possibility - and related to conditions that favor sparse encoding - is that HZE particle exposure increases dentate gyrus neurogenesis. In young adult rodents, inducible increase in hippocampal neurogenesis improves pattern separation, while inducible decrease in neurogenesis impairs pattern separation (58, 80, 88, 134). However, here we show that improved pattern separation is not correlated with the number of new hippocampal neurons. This adds to the growing evidence that the number of new neurons does not always predict pattern separation performance, particularly in older rodents (135–137). In fact, decreased neurogenesis is proposed to diminish sensitivity to memory interference and thus improve performance in certain memory tasks (137–139). Computational models support that decreased neurogenesis may enhance sparse encoding (140, 141), which as mentioned above may explain why we see improved pattern separation yet other groups see decreased performance in their behavioral tests.

The disconnect shown here between pattern separation and hippocampal neurogenesis raises interesting future directions. Although historically tied to learning and memory, hippocampal neurogenesis also plays a role in forgetting (142, 143) with high levels of hippocampal neurogenesis facilitating the forgetting of prior memories, resulting in greater cognitive flexibility (142–144). In converse, lower levels of hippocampal neurogenesis - as seen with age - facilitate the persistence of prior memories, more interference with new memory formation, and thus less cognitive flexibility (143, 144). As here we show irradiated mice have decreased neurogenesis relative to control mice, it is possible irradiated mice have consequently decreased forgetting (greater memory persistence) and also experience more proactive interference from past memories and would have less cognitive flexibility. Rodent cognitive flexibility can be directly tested using a reversal learning paradigm similar to the PD/Reversal learning task presented here. However, this task does not test rodent memory retention, and as we have shown, this relatively simplistic reversal learning is not affected by HZE radiation exposure. If the PD memory load were to be increased - for example, by training with more pairs of images - the rodent’s ability to then perform reversal with this larger number of stimuli would provide a more robust interrogation of cognitive flexibility. Alternatively, future experiments can hone in on dentate gyrus-specific cognitive flexibility via assessed LD reversal (54, 62, 88), which contrasts with the PD reversal reliance on non-dentate gyrus brain regions (primarily PFC, perirhinal cortex, striatal circuits). Specifically, a challenging LD within-session reversal test would provide clarity as to whether IRR mice have decreased dentate gyrus specific-cognitive flexibility relative to controls (62). Finally, future experiments could probe the influence of HZE particle exposure on the converse of pattern separation: pattern completion (i.e. formation of an accurate generalization of partial sensory input) (145–147). Pattern separation and pattern completion abilities have a reciprocal relationship in mice and aged humans (134, 145–147). As we show HZE particle exposure improves pattern separation (fine detail discrimination) and may increase proactive interference (given the decreased neurogenesis), it is possible irradiated mice have improved pattern separation yet worse pattern completion ability. If that were true, we could then further explore the possibility that the functional switch from pattern completion to pattern separation is driven in part by a slowing of the development of adult-generated neurons (134, 148). However, pattern completion relies on memory recall (134), which is assessed in our PAL paradigm (56) and normal in our irradiated mice.

In conclusion, it is understandable that HZE particle exposure is presumed to have a negative influence on some lower and high-level cognitive functions, as many studies support this conclusion (21, 23, 44, 149–151). However, our study shows this is not universally true. Mature mice exposed to two different HZE particles perform similarly to control mice on many high-level cognitive tasks, reflecting the functional integrity of key neural circuits (i.e. PFC-perirhinal cortex-striatum, dorsal striatum, posterior cingulate cortex, hippocampus). Strikingly, irradiated mice actually perform better than control mice in both appetitive and aversive pattern separation tasks. Whether this HZE exposure-induced dentate gyrus-selective functional enhancement is compensation to earlier irradiation-induced neuromorphological changes (152) remains to be tested. However, our work urges revisitation of the generally-accepted conclusion that space radiation is detrimental to cognition.

## MATERIALS AND METHODS

### Animals

Animal procedures and husbandry were in accordance with the National Institutes of Health Guide for the Care and Use of Laboratory Animals, and performed in IACUC-approved facilities at UT Southwestern Medical Center (UTSW, Dallas TX; AAALAC Accreditation #000673, PHS Animal Welfare Assurance D16-00296, Office of Laboratory Animal Welfare [OLAW] A3472-01), Children’s Hospital of Philadelphia (CHOP, Philadelphia, PA; AAALAC Accreditation #000427, PHS Animal Welfare Assurance D16-00280 [OLAW A3442-01]) and Brookhaven National Laboratories (BNL, Upton NY; AAALAC Accreditation #000048, PHS Animal Welfare Assurance D16-00067 [OLAW A3106-01]). 2-month(mon)-old male C57BL/6J mice (Jackson Laboratories, stock #000664) were housed at UTSW and shipped to BNL for irradiation at 6 mon of age. During shipping and housing at BNL, mice were provided Shepherd Shacks (Bio-Serv). Mice were housed at UTSW or BNL (3-4/cage, light on 06:00, lights off 18:00, UTSW: room temperature 68-79°F, room humidity 30-70%, BNL: room temperature 70-74°F and room humidity 30-70%). At both facilities, food and water were provided *ad libitum*.

### Particle irradiation (IRR)

Mice received whole body HZE (^56^Fe: 600 MeV/n, LET 174 KeV/u, Figs. 2**-**5, 7. Figs. S1-3; or ^28^Si: 275 MeV/n, LET 72 KeV/u, Fig. 6) particle radiation at BNL’s NASA Space Radiation Laboratory (NSRL) during NSRL campaigns and 12C, 13A, 13B, 16B, and 18A. The ^56^Fe and ^28^Si ion beams were produced by the AGS Booster Accelerator at BNL and transferred to the experimental beam line in the NSRL. Delivered doses were ±0.5% of the requested value. All mice - regardless of whether control (Sham) or experimental - were placed for 15 minutes (min) in modified clear polystyrene cubes (AMAC Plastics, Cat #100C, W 5.8 x L 5.8 x H 10.6 cm; modified with ten 5-mm air holes). For ^56^Fe experiments, mice received Sham IRR (placed in cubes Monday, Wednesday, Friday, but received no IRR) or either Fractionated (Frac) 20 cGy ^56^Fe (600 MeV/n, LET 174 KeV/μ, Dose rate 20 cGy/min; placed in cubes and received 6.7 cGy on Monday, Wednesday, and Friday), or Non-Fractionated (Non-Frac) 20 cGy ^56^Fe (placed in cubes Monday, Wednesday, and Friday but received 20 cGy only on Friday). For ^28^Si IRR, mice received Sham IRR (placed in cubes, but received no IRR) or a single exposure of either 20 cGy or 100 cGy ^28^Si (275 MeV/n, LET 72 KeV/μ, Dose rate 20 cGy/min or 100 cGy/min). Post-IRR, mice were returned to UTSW or CHOP and housed in quarantine for 1-2 mon prior to initiation of behavior testing. Body weights (Fig. S1A) were taken multiple times: prior to irradiation, at irradiation, and at least monthly post-IRR until collection of brain tissue.

### Overview of behavioral testing

All mice began behavior testing 1-2-mon post-IRR. Parallel groups of mice were tested for appetitive touchscreen behavioral tests (operant touchscreen platform: touchscreen training; Pairwise Discrimination, PD; PD reversal; Location Discrimination, LD; different paired associates learning, PAL; Visuomotor Conditional Learning, VMCL) vs. aversive behavioral tests (contextual fear conditioning, CFC; contextual discrimination fear conditioning, CDFC). Subsets of mice were also tested for general activity (locomotor, LM), anxiety (dark/light box test, D/L) and pain sensitivity (pain threshold, PT), methods for which are provided in **Supplementary Methods.**

### Appetitive Behavior Testing

The touchscreen platform used was Model 80614 made by Lafayette Instruments (Lafayette, IN). Software used for the touchscreen system was ABET II (Lafayette Instruments, Cat. #89505), and individual ABET programs for specific touchscreen training and testing sessions are listed below or in **Supplementary Methods.** Sham and IRR mice were trained on an operant touchscreen platform (TS training), an overview of which is provided below. Additional touchscreen methods are provided in **Supplementary Methods.**

#### Food exposure/restriction

Three days prior to touchscreen training, each cage of Sham or IRR behaviorally-naive, group-housed mice received daily access to Strawberry Ensure (Strawberry Ensure, Abbott Laboratories, Chicago, IL) in a volume sufficient to cover the bottom of a 2” plastic petri dish. TS training and testing occurred Monday through Friday during the light cycle. Mice were maintained on a food-restricted diet (**Supplementary Methods**).

#### Touchscreen training (Abet II software, Cat. #89505)

General touchscreen training (Fig. 2A, Fig.3A) consisted of 5 steps: Habituation (Hab), Initial Touch (IT), Must Touch (MT), Must Initiate (MI), and Punish Incorrect (PI) (**Supplementary Methods**). During general touchscreen training, either a two-window (2×1; Fig. 2A) or three-window (3×1; Fig. 3A) mask was used. Training of Hab, IT, MT, MI was considered complete when mouse finished 25 trials. Latency (days) to complete each training step is reported in Fig. 2A and Fig. 3A. Criteria of PI is 25 trials in 30 min at >76 % accuracy (day 1) and >80% accuracy (day 2) over two consecutive days.

#### Pairwise Discrimination (PD)/Reversal Testing (ABET II software, Cat. #89540)

After training on the touchscreen platform (above, **Supplementary Methods**), mice went through PD/PD Reversal tests (Fig. 2). For PD, two images from the image bank that the mice had never seen before were simultaneously presented on the screen (i.e. plane vs. spider). Only one of these stimulus images was rewarded (S+), and the image that was rewarded was counterbalanced within each group of mice. After the mouse initiated the trial, the rewarded image was presented on either the left or right side of the screen. The presentation side was pseudo-randomly selected such that the S+ was not presented on the same side more than 3 times in a row. An incorrect choice led to a correction trial, and the mouse had to repeat the trial until it correctly selected the rewarded image displayed in the same location. The correction trial was not counted towards the final percent of trials correct. For Reversal testing, the S+ and S- were switched so that the previously-rewarded S+ stimulus image was now no longer rewarded. The mouse performed PD or Reversal testing until it was able to complete 24 trials in 30 min at >76% accuracy (day 1) and >80% (day 2) for 2 days in a row. For PD and PD reversal data, day 1 and 6 and the last day or day 1, 8, 12, and last day were reported, respectively. Day to completion indicated average of days to reach criteria. Distribution of proportion of subjects which reach criteria was plotted over all testing days. Session length (seconds [s]), percent (%) correct, the number of errors (number of correction trials) are also reported.

#### Paired associates learning (PAL, ABET II software, Cat. #89541)

Once mice achieved all five stages of general touchscreen training using the three-window mask (Fig. 1B, 3A), they began training in and assessment on object-location different paired-associates learning (PAL). There were three possible stimulus ‘objects’ (images of a flower, plane, or spider) and three possible positions on the screen (left, middle, or right) (Fig. 3B). All ‘objects’ had a correct ‘location’ that was unique to them. Two stimuli were displayed at the same time during a trial. One was in the correct location (S+) and the other was in the incorrect location (S-), and whether a stimulus was correct was determined by the location in which it was presented (e.g. flower/left; plane/middle; spider/right). If the mouse nose-poked the incorrect stimulus, no reward was delivered and a 5s time-out followed before the mouse was given the opportunity to complete a correction trial. Correction trials continued until the correct stimulus was chosen. A correction trial (the number of errors) consisted of representation of the stimulus array in the same location configuration. Correction trials were not included in the percent correct. Each session was complete when the mouse performed 25 trials or 30 min had elapsed. PAL lasted for 29 days, and measures reported include session length, completed trial number, percent correct, and number of errors.

#### Visuomotor Conditional Learning (VMCL, ABET software, Cat #89542)

For VMCL, mice received one additional training step, termed VMCL train, prior to the VMCL test.

#### VMCL train

VMCL train is designed to teach the mouse to touch two images on the screen in a specific order and in rapid succession. The first touch must be to an image presented in the center of the screen, and the second touch must be to an image presented either on the left or right of the screen. Specifically, after trial initiation, the mouse must touch a center white square (200 x 200 pixels), which then disappears after touch. A second white square immediately appears on either the left or right side of the screen in a pseudorandom style, such that a square is located on each side 5 out of 10 times, but not more than 3 times in a row. If the mouse selects the location with the second white square, a reward is provided, and a 20s inter-trial interval starts. However, if the mouse selects the location without a square, then the second stimulus is removed, and the house light illuminates for 5 s to indicate a timeout period which must conclude prior to the 20-s inter-trial interval. Then the mouse is presented with a correction trial which must be completed prior to a new set of locations being displayed. VMCL train is complete when the mouse completes 2 consecutive days of 25 trials in 30 min with >75% correct. Session length (presented in s), trial number, percent correct, and number of errors are reported on the first and the last day of VMCL train. Day to completion indicates average of days to reach criteria. Distribution of proportion of subjects which reach criteria is plotted over entire train days.

#### VMCL test

Mice are provided with a center black-and-white image (spikes or horizontal bars, Fig. 3G, GH). Once touched, the center image disappears and white squares appear on both the right and left of the screen. For this task, the center image of the spikes signals that the rodent should touch the right square, while the center image of the horizontal bars signals that the rodents should touch the left square. The two center images are presented pseudorandomly for an equal number of times, and the mice have 2 s to touch the white square on the right or left side of the central image. If they fail to touch the white square within 2 s, a timeout period begins. The same timeout and inter-trial-intervals are used for VMCL testing as were used for VMCL train. As with VMCL train, VMCL test correction trials are used to protect against side bias. VMCL testing is complete when the mouse completes 2 consecutive days of 25 trials in 30 min with ≥76% correct (day 1) and ≥80% correct (day 2) in a row. Session length (presented in s), trial number, percent correct, percent missed, and number of errors are reported for Day 1 and 8 and the last day of VMCL test. Day to completion indicates average of days to reach criteria. Distribution of proportion of subjects which reach criteria is plotted over all VMCL test days.

#### Location Discrimination (LD; ABET2 software, Cat #89546-6)

For LD, mice receive one additional training step, termed LD1-choice, prior to the actual 2-choice LD test (LD2).

#### LD train

Mice initiate the trial, which leads to the display of two identical white squares (25 x 25 pixels, Fig. 4A) presented with two black squares between them, a separation which is termed “intermediate” in difficulty (8^th^ and 11^th^ windows in 6 × 2 high grid-bottom row). One of the locations of the squares is rewarded (L+) and the other is not, and the L+ location (left or right) is counterbalanced within-group. On subsequent days, the rewarded square location is switched (becomes L-), then L+, then L-, etc. A daily LD train session is complete once the mouse touches either L+ or L- 25 times or when 30 min has passed. Once the animal reaches 25 trials in 30 min for 2 consecutive days (irrespective of accuracy), the mouse advanced to the LD 2-choice random test. Session length and percent correct on the last day of LD train are reported. Days to completion indicates average of days to reach criteria and distribution of proportion of subjects which reach criteria is plotted over entire training days.

#### LD 2 Random choice EH (referred to hereafter as ‘LD test’)

Mice initiate the trial, which leads to the display of two identical white squares, either with four black squares between them (“large” separation, two at maximum separation (7^th^ and 12^th^ windows in 6 x 2 high grid-bottom row) or directly next to each other (“small” separation, two at minimum separation (9^th^ and 10^th^ windows in the Bussey Mouse Operant Mode 6 x 2 high grid-bottom row; Fig. 4F). Like the LD 1 train, only one of the square locations (right-most or left-most) is rewarded (L+, same side for both large and small separations, and counterbalanced within-groups). The rewarded square location is switched the following day, and the location continues to alternate daily throughout testing. Each day, the separation (large vs. small) is pseudorandomly displayed (same separation shown no more than 3 consecutive times). LD testing is complete when the mouse completes 45 trials in 30 min regardless of accuracy. Session metrics reported are length, percent correct, number of completed trials, number of blank touches, reward collection latency (time between reward presentation and the first head entry into the reward port), and correct/incorrect image response latency (latency from correct/incorrect image response). For analysis of performance in 10-trial Blocks (1^st^ 10-trial Block: 1-10 trials, 2^nd^ 10-trial Block: 11-20 trials, 3^rd^ 10-trial Block: 21-30 trials, 4^th^ 10-trial Block: 31-40 trials) on the last day, metrics reported are duration, percent correct, blank touch, and left and right touch during inter-trial-interval. Days to completion indicates average of days to reach criteria and distribution of proportion of subjects which reach criteria is plotted over all LD test days.

#### Aversive Behavior Testing

CDFC overview is provided below. See Fig. S2 **and** S3 and **Supplementary Methods** for additional CDFC information, and for detailed information about CFC. *Contextual Discrimination Fear Conditioning (CDFC).* A modified CDFC behavioral paradigm was utilized in which mice were exposed daily to two contexts (Context A and B) that shared similarities (including a floor pattern, a high-salience contextual feature (153)) but had distinct visual and olfactory features (Fig. S2)(82, 134), and were paired with distinct handling approaches (**Supplementary Methods**). Importantly, Context A was always paired with a foot shock, while Context B was never paired with a foot shock, as described below.

Over the course of 16 days, mice were exposed daily to both Context A and Context B. The order of exposure to Context A and B alternated between days (BAABABBABAABABBA) such that on days 2, 3, 5, 8, 10, 11, 13, and 16 mice were exposed to Context A first and Context B second (Fig. S2). For CDFC data analysis, the percent freezing in Context A and Context B were measured each day, and data from each treatment group were collapsed and averaged across every two days, referred to as Blocks. Therefore, data were analyzed as 8 Blocks (16 testing days) such that the grouping of days into Blocks was as follows: [BA AB] [AB BA] [BA AB] [AB BA] etc. However, since Day 1 of exposure includes data from mice prior to their first tone/shock pairing and therefore their response does not reflect a learned association, Block 1 (Days 1-2) was removed from analysis. Percent of time freezing was measured using linear analysis. The threshold for freezing was 20 arbitrary units detected using the proprietary Med Associates Software. Additional analysis parameters include bout length (0.5 s) and frames/s (30).

### Tissue Collection

After completion of behavioral tests, mice underwent intracardial perfusion and fixation as previously described (70, 154, 155). ^56^Fe IRR mice were perfused 4-6-mon post-IRR (10 to 12 mon of age) and ^28^Si IRR mice were perfused 6-mon post-IRR (14 mon of age). Briefly, mice were anesthetized with chloral hydrate (Sigma-Aldrich cat. #C8383, 400 mg/kg, stock solution 400 mg/ml made in 0.9 % NaCl solution, i.p.) and exsanguinated intracardially with 0.1M PBS (7 ml/min, 6 min) and followed by perfusion intracardially with 4 % paraformaldehyde in 0.1M PBS (7 ml/min, 15 min). As stress can influence neurogenesis and thus doublecortin-immunoreactive (DCX+) cell number, steps were taken to minimize potential stress differences among mice in the same cage: each cage was gently removed from the housing room and brought to the adjacent procedure room immediately prior to anesthesia; mouse cage transfer was performed by a researcher with clean personal protective equipment; and all mice in a cage were anesthetized within 3 min and began exsanguination within 5 min of being brought into the procedure room. With these and other steps, we have found neurogenesis levels in mice can be reliably and accurately evaluated. Brains were harvested and placed in 4% paraformaldehyde at room temperature for 2 days, transferred to cryoprotectant (30% sucrose in 0.1 M PBS and 0.1% NaN_3_) and stored at 4°C until sectioning. Brains were coronally sectioned on a freezing microtome (Leica), with 30 μm sections collected in serial sets through the entire anterior-posterior length of the hippocampus (distance range from Bregma: −0.82 to −4.24 μm)(156). These eight serial sets of sections (section sampling fraction, ⅛) were stored in 0.1% NaN_3_ in 1x PBS (Fisher Scientific; Pittsburgh, PA) at 4°C until processed.

### Immunohistochemistry

Immunohistochemistry was performed as previously described (70–72). Briefly, one complete set of coronal sections from a 1:n series (1:8 or 1:9) was mounted onto glass slides (Superfrost/Plus, Fisher) in rostral to caudal order and allowed to dry. To visualize DCX+ cells using 3’3-diaminobenzidine (DAB), slide-mounted sections were treated for antigen retrieval (0.01M citric acid in MQH_2_O, pH 6.0, 95°C, 15 min) and quenching of endogenous peroxidases (0.3% hydrogen peroxide in 1xPBS, 30 min). Non-specific staining was blocked by incubation in 3% normal donkey serum (NDS) and 0.1% Triton X-100 in 1xPBS for 60 min. Sections were then incubated in goat-anti-DCX primary antibody (1:500, Santa Cruz) overnight at room temperature in 3% NDS, 0.1% Tween-20 in 1xPBS. The following day, sections were incubated for 60 min with biotinylated donkey anti-goat antibody (1:200, Jackson ImmunoResearch) in 1.5% normal donkey serum in 1xPBS followed by rinses. A 60-min incubation in avidin-biotin complex (ABC Elite, 1:50, Vector Laboratories) was then performed, followed by visualization of immunoreactive cells using DAB (Thermo Scientific Pierce) and Nuclear Fast Red counterstaining (Vector Laboratories). Tissue was then dehydrated with a series of increasing ethanol concentrations and defatted section (Citrosolv) were cover slipped with DPX Mountant (Sigma-Aldrich).

### Stereological Cell Quantification

Unbiased analysis of DCX+ cell number was performed via stereologic quantification on a BX51 System Microscope (Olympus America, Center Valley, PA, USA) as previously described (70–72). DCX+ cells in the subgranular zone and granular cell layer of the hippocampal dentate gyrus were visualized with a 40X, 0.63 NA oil-immersion objective and quantified with the formula:

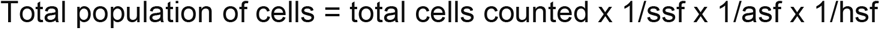

where ssf is the section sampling fraction (DCX: e.g. ⅛), asf is the area sampling fraction (1 for these rare populations of cells; thus, all cells were counted in ⅛ of the sections), hsf is the height sampling fraction (1 given the minimal effect edge artifacts have in counting soma <10μm with ssf ⅛). As both hemispheres were counted for DCX, the resulting formula was:

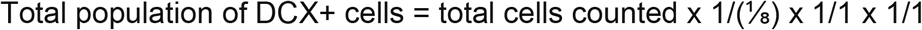

### Statistical Analyses

Data are reported as mean ± s.e.m. Testing of data assumptions (for example, normal distribution, similar variation between control and experimental groups, etc.) and statistical analyses were performed in GraphPad Prism (ver. 8.2.0). Statistical approaches and results are provided in Table S1 for main figures and in Table S2 for supplementary figures, and statistical analysis summaries are provided in the figure legends. Analyses with two groups were performed using an unpaired, two-tailed Student’s t-test, and significance is indicated by asterisks (e.g., *p < 0.05, **p < 0.01, ***p < 0.001). Analyses with more than two groups and one variable were performed using one-way ANOVA and Bonferroni post hoc test; post hoc significance is indicated by asterisks (e.g., *p < 0.05, **p < 0.01, ***p < 0.001). Analyses with more than two variables were performed using two-way ANOVA with Bonferroni post hoc test; repeated measures (RM) were used where appropriate, as indicated in figure legends and Table S1. Two-way ANOVA post hoc significance is indicated by lowercase letters (e.g., *a, b, c*: p < 0.05; *a’, b’, c’*: p < 0.01; *a’’, b’’, c’’*: p < 0.001). For the distribution of subjects reaching criteria between control and experimental groups, the Mantel-Cox test was used, and significance was defined as *p < 0.05. For behavioral studies, mice were randomly assigned to groups. Additionally, investigators were blinded to the treatment group until all data had been collected. Sample sizes were pre-determined via power analysis and confirmed on the basis of extensive laboratory experience and consultation with CHOP and PennMed statisticians.

### Figure Preparation

For graphical data, figures for each data set were produced in Prism (GraphPad ver. 8.2.0) and transferred to Illustrator (Adobe Illustrator cc2018 version 22.1) to enable uniform line thickness and figure size. For photomicrographs, immunostained sections were visualized with an epifluorescence microscope (Olympus BX51) with 10x and 40x objectives and images were captured with the Olympus DP Manager Program before being prepared in Adobe Illustrator 2018 (version 22.1).

### Transparency and Reproducibility

Behavioral experiments were performed by researchers blind to treatment (Sham or IRR), which was feasible since such the low doses of space radiation used here do not have gross measurable impact on mouse weight or hair loss. Automated scoring was used for most behavior tests. Touchscreen testing criteria was based on rodent performance, thus avoiding scoring discrepancies among researchers. For immunohistochemical experiments, tissue was coded to obscure treatment information, and codes were not broken until data analyses were complete. After publication, raw data and images will be made available to interested researchers.

## Competing Interests

The authors declare no competing financial interests.

## Author Contributions (based on Project CRediT)

listed via initials and alphabetically by last name

Conceptualization: SGB, BPCC, AJE, MJL, CWW, SY

Methodology: SGB, BPCC, AJE, MJL, IS, FT, CWW, SY

Software: Not applicable

Validation: MJL, RPR, IS, CWW, SY

Formal Analysis: SGB, AJE, NI, MJL, DRR, PDR, RLR, IS, HYS, FT, CWW, SY

Investigation: SGB, NAD, AJE, NI, MJL, SM, GP, RLR, DRR, PDR, HYS, CWW

Resources: SGB, AJE, AMS

Data Curation: AJE, MJL, RPR, CWW, SY

Writing, original draft: AJE, CWW, SY

Writing, review and editing: SGB, AJE, IS, CWW, SY

Visualization: AJE, RPR, CWW, SY

Supervision: AJE, AMS, SY

Projection Administration: AJE, AMS

Funding Acquisition: BCC, AJE, SY

## Funding and Acknowledgements

Research supported by NASA grants NNX07AP84G (to BPC and AJE), NNX12AB55G (to AJE and BPC), and NNX15AE09G (to AJE) and NIH grants to AJE DA007290, DA023555, and DA016765. CWW was supported by an NIH Institutional Training grant (DA007290, PI: AJE, David W. Self), and SY was supported by an NIH Institutional Training Grant (MH076690, PI: CA Tamminga), a PENN McCabe award, and an IBRO travel grant. We thank members of the Eisch, Chen, and Stowe Laboratories for technical support and helpful conversations such as Lyles Clark, Fred Kiffer, Guillermo Palchik, Shibani Mukherjee, Vanessa Torres, Angela K. Walker, and Kielen Zuurbier. We thank members of the Brookhaven National Laboratory staff including Adam Rusek (Physics team leader), MaryAnn Petry (animal support director), Peter Guida (organization and technical support director) as well as all of their team members who help make our experiments possible.

## Datasets

Raw data are made available to researchers on written request.

## Ethics

Human subjects: No

Animal subjects: Yes

Ethics statement: All experiments were conducted in accordance with the regulations of the USDA and the IACUC at UTSW, CHOP, and BNL.

## Dual-use research

Not applicable.

## Permissions

This manuscript represents original work, and is not a reproduction or modification of any part of an article that has been previously published or submitted to another journal.

## SUPPORTING INFORMATION APPENDIX

### SUPPLEMENTARY MATERIAL AND METHODS

#### Locomotor Activity (LM)

Within 2-months (mon) post-IRR (^56^Fe experiments: 59-days post-IRR; ^28^Si experiments: 49-days post-IRR), mice underwent a single locomotor activity recording session from 5pm-9am under red light. After 1-hour (hr) acclimation to the testing suite, group-housed mice were individually placed into clean standard cages and were given *ad libitum* food and water. Beam breaks were recorded over 16 hr using the Photobeam Activity System-Home Cage (San Diego Instruments; San Diego, CA). Data were collapsed into 30-minute (min) bins across the 16-hr session, and are presented as number of beam breaks. At the completion of recording, mice were placed back to their original group-housed cage and returned to their normal housing room.

#### Dark/Light test (D/L)

The apparatus consisted of a polypropylene cage (L 44 x W 21 x H 21 cm) unequally divided (⅔ and ⅓) into two chambers. The large chamber was white and brightly-illuminated by two 20-W fluorescent lights (1388 lux at cage floor), while the small chamber was dark (not illuminated). Initially the mouse was placed in the dark side for 2 min, after which the door between the two chambers is opened and the transitions of the mouse between the two chambers and time in each chamber was detected for 10 min by seven photocells. The time spent in the brightly-lit side and latency to enter the brightly-lit side were measured by an automated system (Med Associates).

#### Pain Threshold (PT)

Mice were individually placed into boxes equipped with a metal grid floor connected to a scrambled shock generator (Med Associates Inc., St. Albans, VT). After ∼1 min, mice received a series of foot shocks (each 2-second [s] duration) with increasing intensity. The initial shock intensity was 0.05 mA, and the amplitude was increased by 0.05 mA for each consecutive foot shock with a 15-s intershock interval. The first shock intensity at which each animal displayed each behavior (flinch, vocalization, or jump) is reported. Once the animal displayed all three behaviors, it was removed from the chamber.

#### Contextual Discrimination Fear Conditioning (CDFC)

CDFC paradigm and chambers are shown and described in Figure S2. *“*Context A” consisted of a standard fear conditioning chamber (Med Associates) outfitted with a grid floor and white overhead house light, was scented with vanilla, and was paired with a shock. “Context B” consisted of a standard fear conditioning chamber with a grid floor, but with a near-infrared light and a black A-frame insert, was scented with mint, and was not paired with a shock. There were other subtle differences between the contexts. For example, prior to placement into Context A, mice were individually placed into a transfer cage (a standard cage with bedding), and then placed by the tail into Chamber A. After exposure to Context A, the mouse was removed and Context A was cleaned with Coverage Plus NPD solution (Steris, Mentor, OH). In contrast, prior to placement into Context B, mice were individually placed into a transfer cage lined with white paper towels, and each mouse was scooped by hand into both the transfer cage and testing chamber. After exposure to Context B, the mouse was removed and Context B was cleaned with 1% acetic acid. Each twice daily exposure over 16 days lasted 4 min 2 s, during which freezing behavior was scored for the first 3 min. Mice in Context A, but not Context B, received a single, mild foot shock (0.25 mA, 2-s duration) after 3 min in the context. Mice then remained in the chambers for one additional minute until the session was complete. The interval between daily exposures to Context A or B was 2-2.5 hrs.

#### Contextual Fear Conditioning (CFC)

CFC paradigm and chambers are shown and described in Figure S3. CFC consisted of two phases: training (Day 1), and testing (Day 2-3). Mice were habituated to the behavior room environment 1 hr each day prior to training and testing sessions. On Day 1, mice were trained to associate a novel context (standard fear conditioning chamber, grid flooring, no odor, house lights on; Med Associates Inc., St. Albans, VT) with a shock. Two minutes after placement in the novel context, an auditory cue was played (80-decibel [dB] white noise, 30-s duration, Med Associates Inc.), which co-terminated with the presentation of a 0.5-mA shock (2-s duration). This cue-shock pairing was repeated twice during Day 1 (5-min training session), with 1 min between the cue-shock presentations. On Day 2, mice underwent context testing: 5 min in the same environment as Day 1 training, but no auditory cue or foot shock presented. On Day 3 (^56^Fe IRR mice) mice underwent auditory cue testing: 6 min in another novel context (plastic flooring, triangular roof, vanilla odor, house lights on). For training and testing sessions, freezing behavior was assessed using VideoFreeze software (Med Associates Inc.), compiled for each phase of each session (e.g. Pre-Cue, During Cue, etc.), and presented as percent percent time freezing for each phase.

#### Touchscreen behavior tests (Abet II software, Cat. #89505)

##### Touchscreen platform and software

The touchscreen platform used was Model 80614 made by Lafayette Instruments (Lafayette, IN). Each operant chamber is encased in a sound-attenuating chamber. Each chamber is trapezoid-shaped, with the widest wall serving as the “touchscreen” (W 238 x H 170 mm) and the opposite and narrowest wall (W 46 mm) containing a motion-sensitive center dispenser (tray) to deliver liquid reward (Strawberry Ensure, Abbott Laboratories, Chicago, IL). Each chamber has two lights (tray light and overhead house light), and is equipped with a speaker (ceiling in each chamber) to play a tone. Aside from initial priming reward used during training, a “reward” is defined as 7uL Ensure delivered to the illuminated tray at the same time as a tone is played. Aside from training sessions, the term “initiate a trial” is defined as the mouse placing its head in the tray when the tray light is illuminated and the tone is played. The two remaining walls of the chamber are infrared-permeable to track rodents during testing. The floor is a perforated metal grid, and the solid roof is hinged for easy placement/removal of the animal. A computer outside of the chamber controls the programs and recording of each session. Mice are tested in their light cycle Monday through Friday until testing was complete. Software used for the Touchscreen System is from ABET II (Lafayette Instruments, Cat. #89505), and individual ABET programs for specific touchscreen training and testing sessions are listed below.

##### Food exposure/restriction

In brief, mouse chow (16% protein 2016 Teklad Global Diet, Envigo, Madison, WI) was removed from each cage at 5 pm the day prior to training or testing. Each cage was given *ad libitum* access to chow for 3 hr (minimum) to 4 hr (maximum) immediately following daily touchscreen training/testing, and from completion of training/testing on Friday until Sunday 5 pm. Mice were weighed each Wednesday to ensure weights >80% initial body weight. While weights below this threshold merited removal of the mouse from the study, zero mice reached this threshold (Fig. S1A).

Touchscreen training (Fig. 2A, 3A) consists of 5 steps, as previously published (54, 56, 109): Habituation, Initial Touch, Must Touch, Must Initiate, and Punish Incorrect. Methods for each step of the touchscreen training are described in turn below.

##### Habituation (Hab)

Mice are placed in touchscreen chamber for 30-min (max) session with the tray light turned on (LED Light, 75.2-lux). For the initial reward in each habituation session, a tone is played (70-dB at 500 Hz, 1000 ms) at the same time as a priming reward (150 uL Ensure) is dispensed to the chamber tray. After the mouse inserted its head and removed its head from tray, the tray light turns off and a 10-s delay begins. At the end of the delay, the tray light is turned on and tone is played again as a standard reward (7uL Ensure) is dispensed. If the mouse’s head remains in the tray at the end of the 10-s delay, an additional 1-s delay is added. Mice complete Habituation training after they collect 25 rewards (25 x 7 ul) within 30 min. Mice that achieve habituation criteria faster than 30 min are removed from the chamber immediately after their 25^th^ reward in order to minimize extinction learning.

##### Initial Touch (IT)

Drawing from a bank of 40 preselected black and white images (240 x 240 pixels), a random image is displayed on the screen in a pseudo-random location such that no image is displayed in that location more than 3 consecutive times. The mouse has 30 s to touch the image (typically with their nose). If the mouse does not touch the image, the image is removed, a reward (7uL Ensure) is delivered into a lit tray, and a tone is played. After the reward is collected, the tray light turns off and a 20-s intertrial interval begins. If the mouse touches the image on the screen while it is displayed, the image is removed and the mouse receives 3 times the normal reward (21 uL Ensure, tray lit, tone played). Mice advance from Initial Touch training after they complete 25 trials (irrespective of reward level received) within 30 min. Mice that achieve Initial Touch criteria faster than 30 min are removed from the chamber immediately after their 25^th^ trial.

##### Must Touch (MT)

Similar to Initial Touch training, a random image is displayed, but now the image remains on the screen until it is touched. If the mouse touches the screen, the mouse receives a reward (7uL Ensure, tray lit, tone played). If the mouse touches the blank screen, there is no response (no reward dispensed, no light in tray, no tone). Mice advance from Must Touch training after they complete 25 trials within 30 min. Mice that achieve Must Touch criteria faster than 30 min are removed from the chamber immediately after their 25^th^ trial.

##### Must Initiate (MI)

Must Initiate training is similar to Must Touch training, but a mouse is now required to initiate the training by placing its head into the already-lit tray. A random image from the image bank will then appear on the screen, and the mouse must touch the image to receive a reward (7uL Ensure, tray lit, tone played). Following the collection of the reward, the mouse must remove its head from the tray and then reinsert its head to initiate the next trial. Mice advance from Must Initiate training after they complete 25 trials within 30 min. Mice that achieve Must Initiate criteria faster than 30 min are removed from the chamber immediately after their 25^th^ trial.

##### Punish Incorrect (PI)

Punish Incorrect training builds on Must Initiate training, but here if a mouse touches a portion of the screen that is blank (does not have an image), the overhead house light turns on and the image disappears from the screen. After a 5-s timeout period, the house light turns off, and the mouse has to initiate a correction trial, where the same image appears in the same location on the screen. The correction trials are repeated until mouse successfully presses the image but are not counted towards the final percent correct criteria. Mice advance from Punish Incorrect training and onto testing after they complete 25 trials within 30 min at ≥76% (≥19 correct) for two consecutive days. Mice that achieve Punish Incorrect criteria faster than 30 min are removed from the chamber immediately after their 25^th^ trial.

---

**Figure S1.**
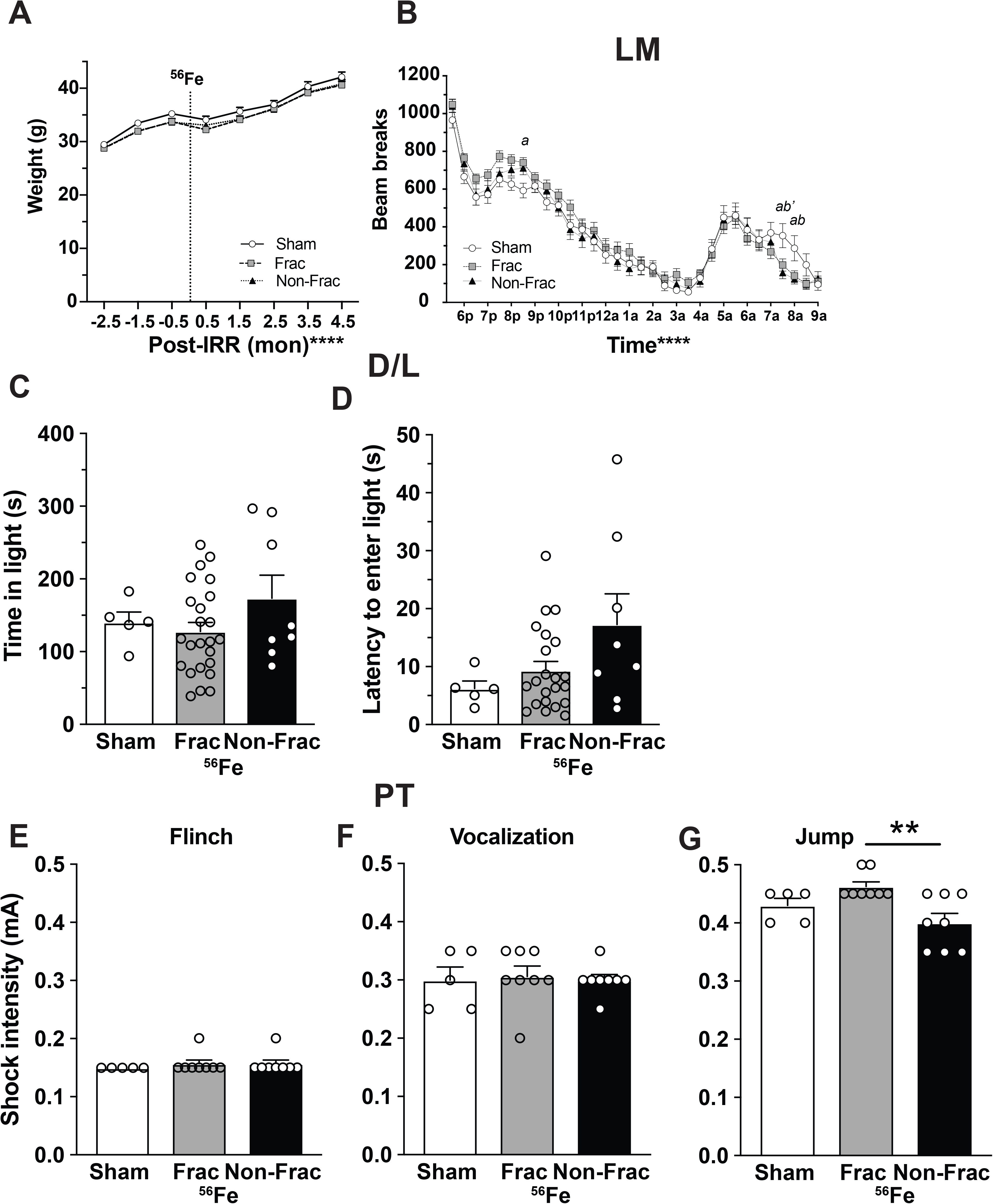
Weights, locomotion, anxiety, and pain threshold are generally unaffected in mice exposed to whole body Frac or Non-Frac ^56^Fe radiation in maturity. **(A)** No gross weight difference was detected before and after radiation in Sham or ^56^Fe groups. **(B-G)** Locomotor activity (LM) measured in 30 minute bins for 16 hrs **(B)**, time spent in light **(C)**, latency to enter light **(D)** in dark/light box (D/L) and measurements for flinch **(E)**, vocalize **(F)**, and jump **(G)** in the pain threshold test (PT) reveal no gross changes after exposure to Sham or ^56^Fe radiation. Mean±SEM. Statistical analysis in **A, B**: Two-way RM measures ANOVA, **** p<0.0001, Bonferroni’s post-hoc analysis, *a* p<0.05 in Sham vs Frac; *b* p<0.05, *b’* p<0.01 in Sham vs Non-Frac. One-way ANOVA in C-G, Bonferroni’s post-hoc analysis. ** p<0.01. a=A.M., Frac=fractionation, months=mon, mA=milliampere, p=P.M., Non-Frac=non-fraction, s=seconds.

**Figure S2.**
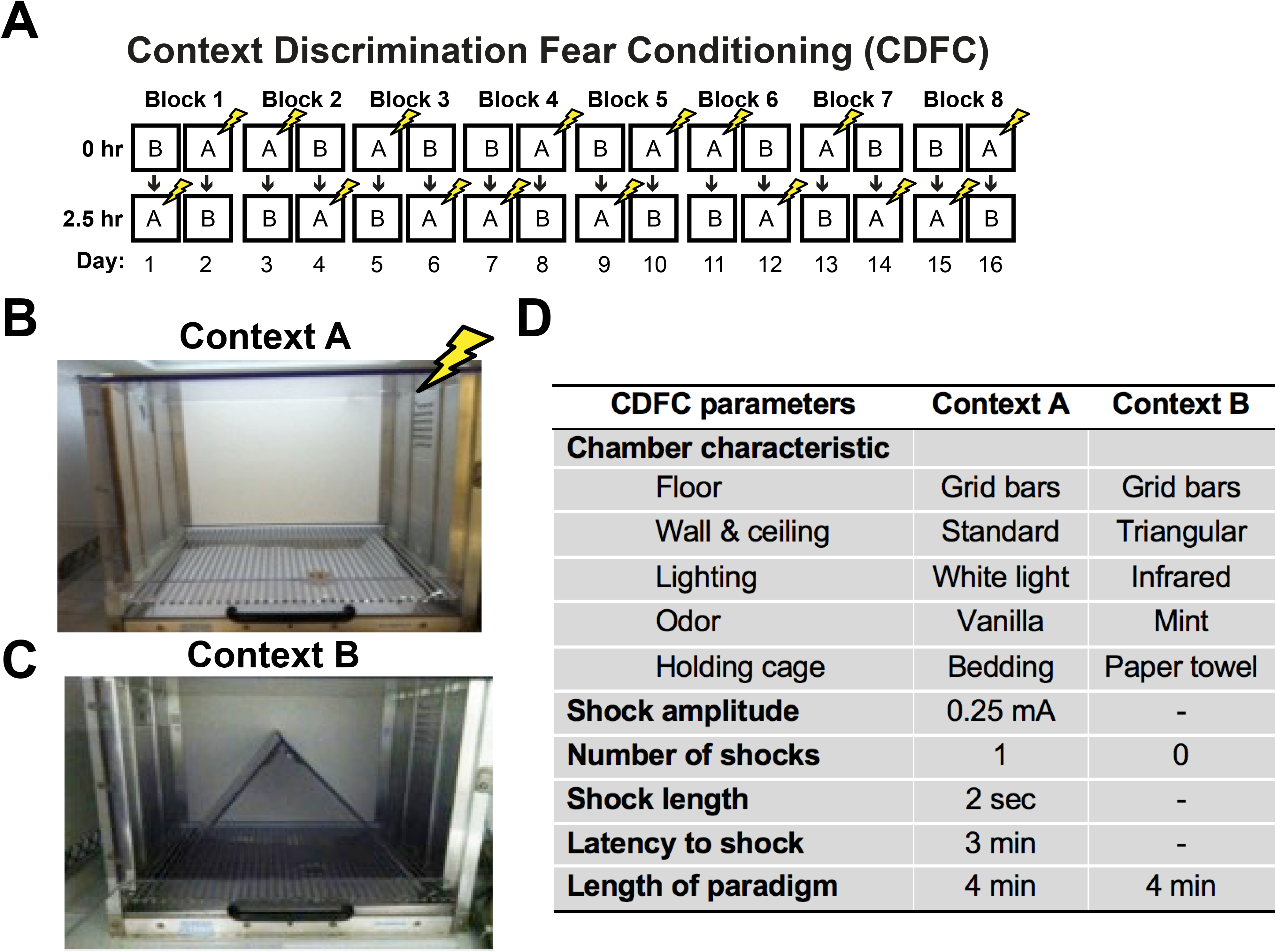
Contextual Discrimination Fear Conditioning (CDFC) paradigm. **(A)** Sixteen-day CDFC paradigm depicting daily, randomized placement into Context A (shock-paired, indicated by yellow lightning bolt) and the contextually-similar Context B (no shock). **(B-C)** Photographs of chamber set up as Context A (**B**, the context paired with mild foot shock) and Context B (**C**, a somewhat distinct context that is never paired with a foot shock). **(D)** Table of parameters of Context A and Context B used for this CDFC paradigm. “-”=not applicable.

**Figure S3.**
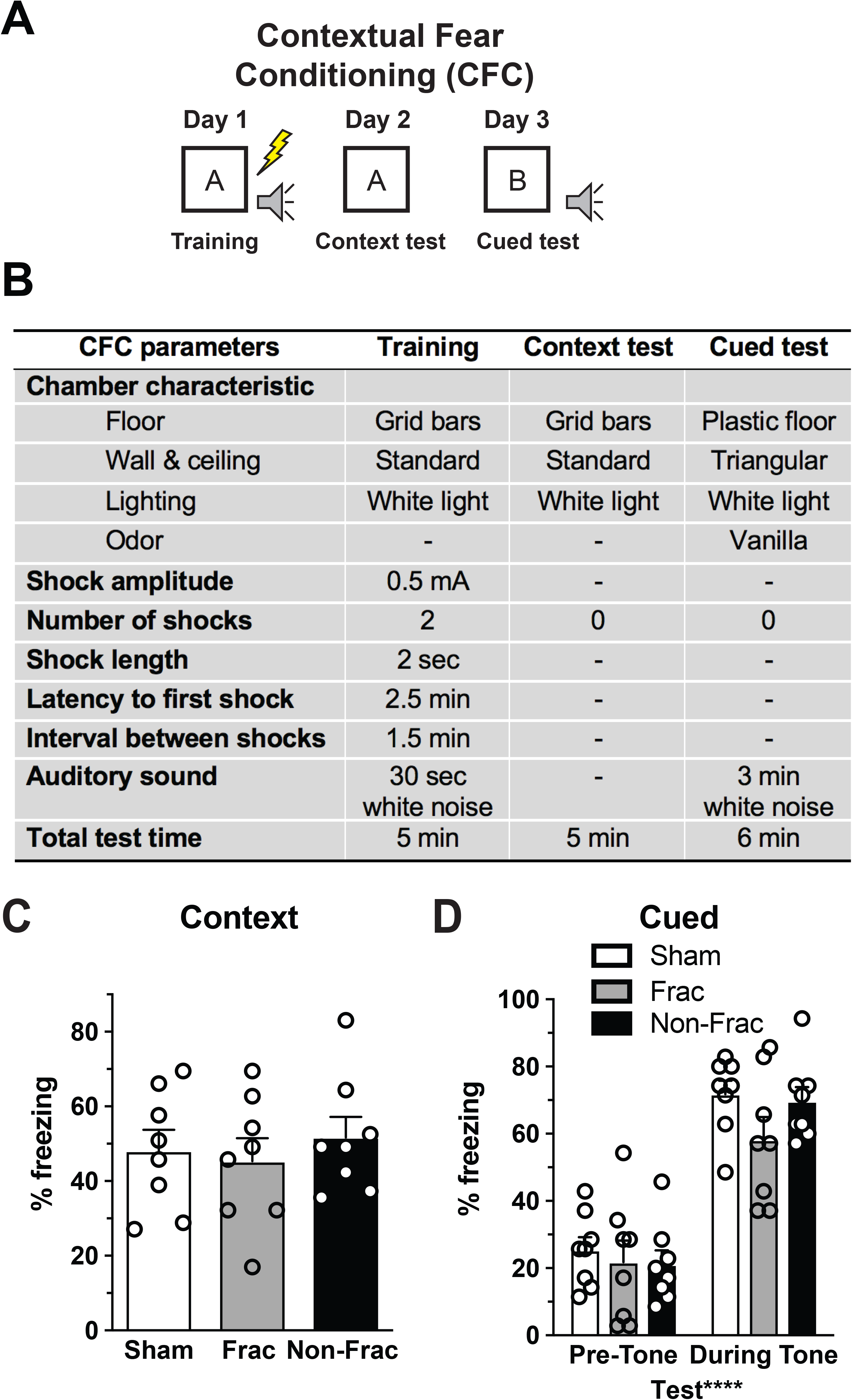
Contextual fear conditioning (CFC) is unaffected in mice exposed to whole body Fractionated (Frac) or Non-Fractionated (Non-Frac) 20 cGy ^56^Fe radiation. **(A)** Three-day CFC paradigm depicting placement (Day 1) into in novel context which is paired with a cue (auditory tone, indicated by grey speaker, is paired with shock, indicated by yellow lightning bolt) followed by testing in the same context (Day 2) and in an additional novel context for cued testing (Day 3). **(B)** Table of parameters of the contexts used for training and testing in this CFC paradigm. **(C-D)** Percent freezing in response to context **(A)** or cue **(B)** in the CFC test reveals a lack of effect with ^56^Fe radiation. Mean±SEM. **(C)** One-way ANOVA, **(D)** Two-way repeated measures ANOVA. ****p>0.0001, “-”=not applicable.

**Table S1.**
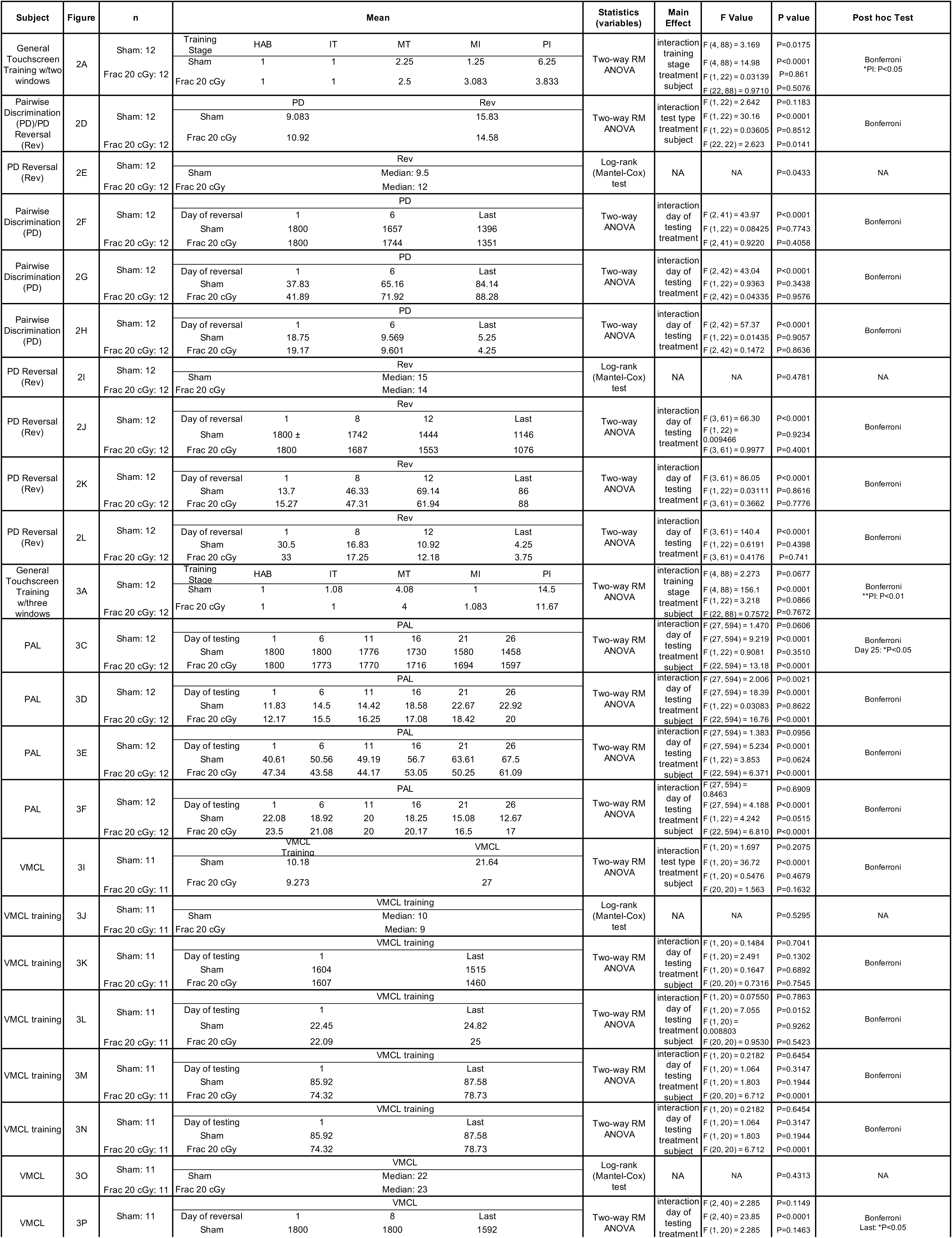
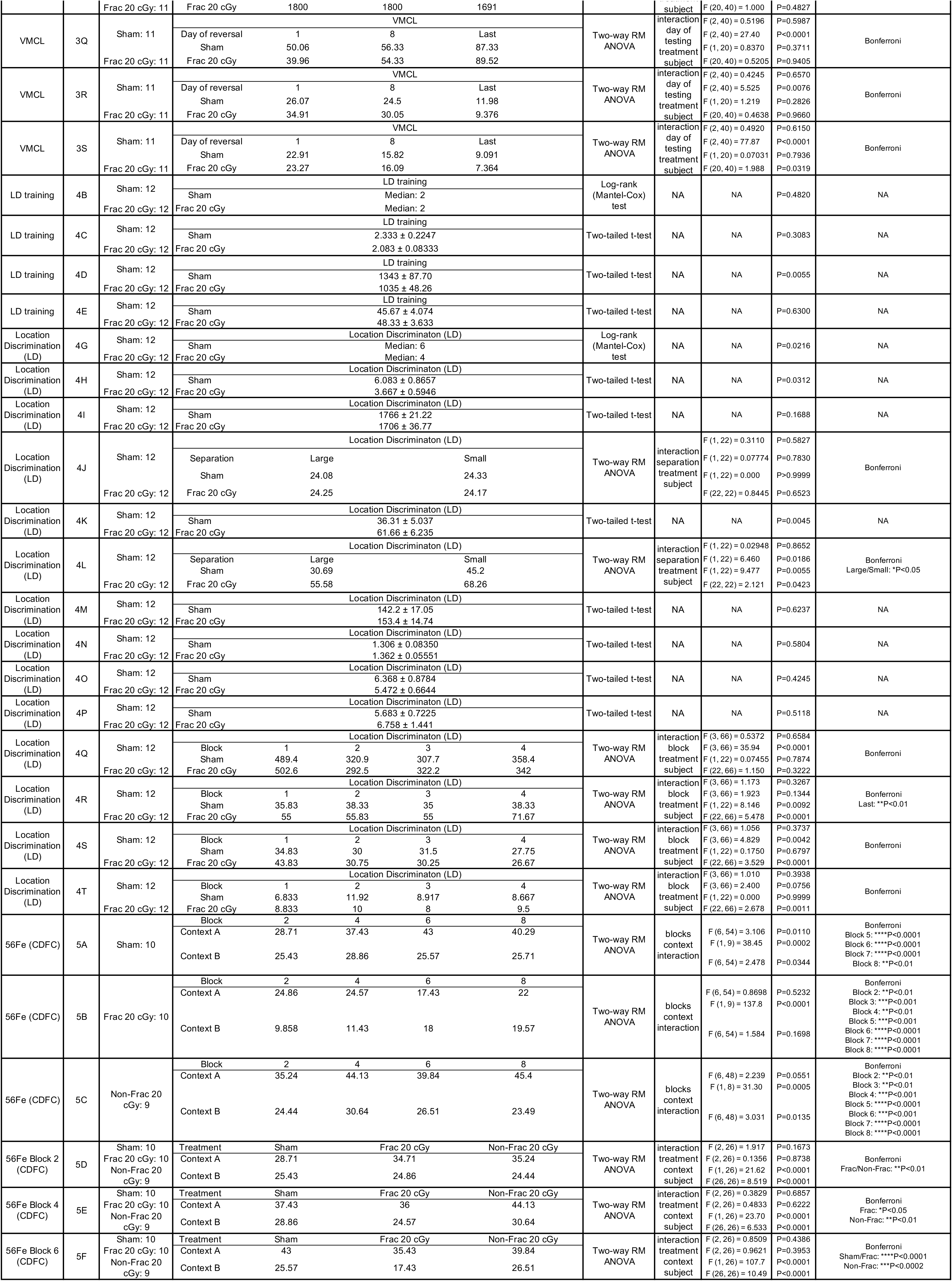
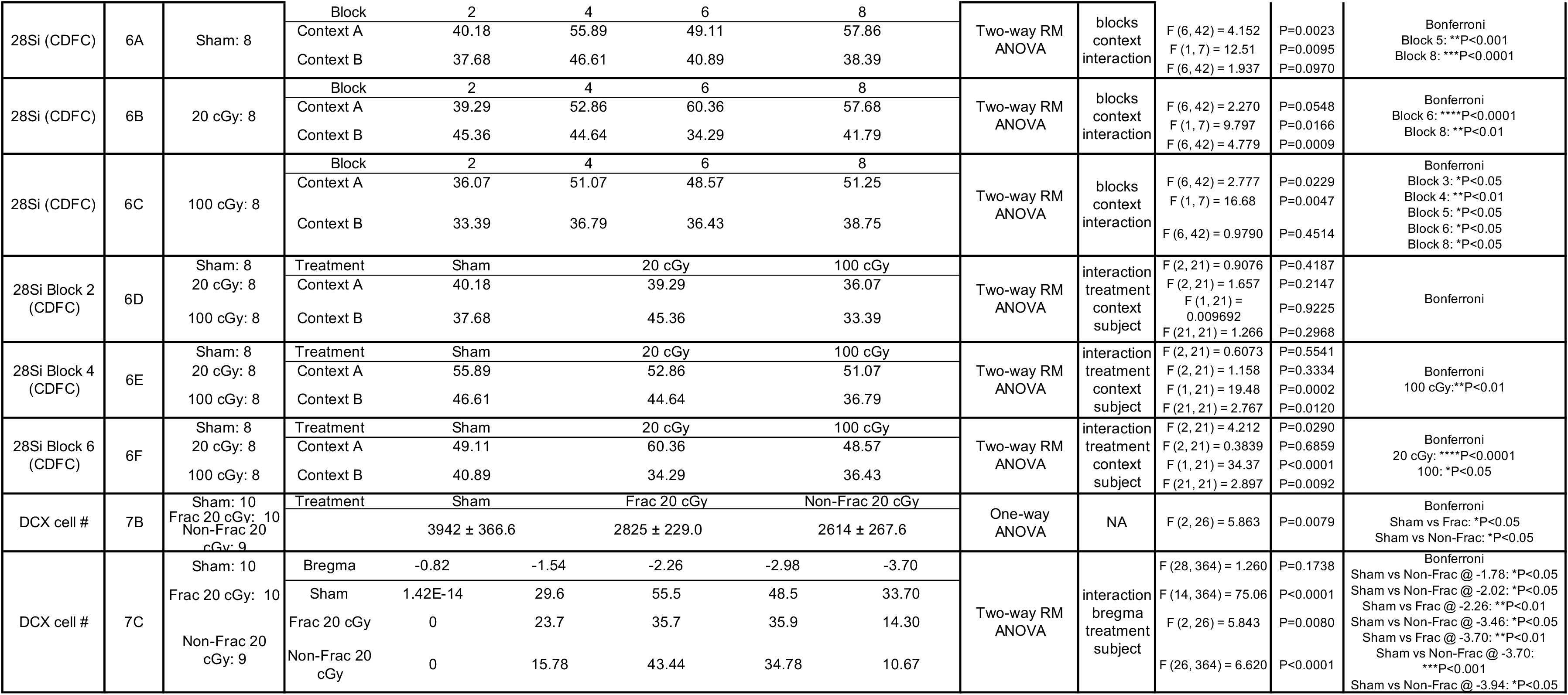
Reporting statistical results of main figures.

**Table S2.**
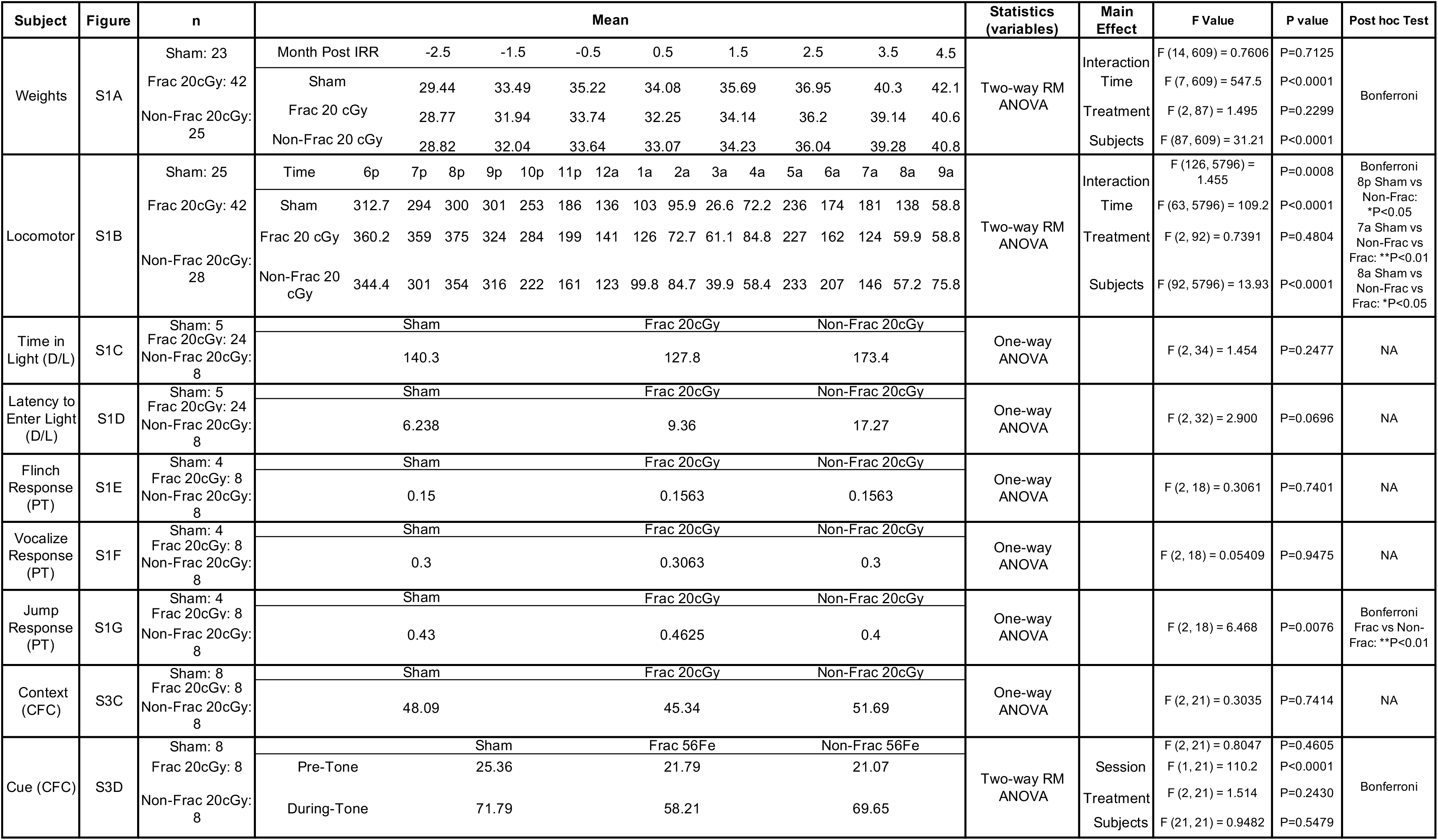
Reporting statistical results of supplementary figures.

